# Identifying states of collateral sensitivity during the evolution of therapeutic resistance in Ewing’s sarcoma

**DOI:** 10.1101/2020.02.11.943936

**Authors:** Jessica A. Scarborough, Erin McClure, Peter Anderson, Andrew Dhawan, Arda Durmaz, Stephen L. Lessnick, Masahiro Hitomi, Jacob G. Scott

## Abstract

Advances in the treatment of Ewing’s sarcoma (EWS) are desperately needed, particularly in the case of metastatic disease. A deeper understanding of collateral sensitivity, where the evolution of therapeutic resistance to one drug aligns with sensitivity to another drug, may improve our ability to effectively target this disease. For the first time in a solid tumor, we produced a temporal collateral sensitivity map that demonstrates the evolution of collateral sensitivity and resistance in EWS. We found that the evolution of collateral resistance was predictable with some drugs, but had significant variation in response to other drugs. Using this map of temporal collateral sensitivity in EWS, we can see that the path towards collateral sensitivity is not always repeatable, nor is there always a clear trajectory towards resistance or sensitivity. Identifying transcriptomic changes that accompany these states of transient collateral sensitivity could improve treatment planning for EWS patients.

## Introduction

Ewing’s sarcoma (EWS) is the second most common primary malignant bone cancer in children (Ries, 1999; Esiashvili et al., 2008). Localized disease has a 50-70% 5-year survival rate, and metastatic disease has a devastating 18-30% 5-year survival rate (Esiashvili et al., 2008; Grier et al., 2003; Hunold et al., 2006). Advances in the treatment of EWS are desperately needed, particularly in the case of metastatic disease. Unfortunately, all recent attempts to improve the chemotherapy regimen for EWS have only yielded modest results for non-metastatic cancer with little-to-no impact on the course of metastatic disease (Grier et al., 2003; Huang and Lucas, 2010). Researchers have tried adding ifosfamide and etoposide to standard EWS chemotherapy, increasing the drug doses administered, and decreasing the interval between doses, all without meaningful improvement to metastatic disease outcomes (Grier et al., 2003; Huang and Lucas, 2010; Womer et al., 2012). Even when treatment is initially successful, EWS often evolves therapeutic resistance, which ultimately leads to disease relapse (Ahmed et al., 2014). A deeper understanding of the evolutionary dynamics at play as EWS develops therapeutic resistance may improve our ability to effectively target this disease (Scott and Marusyk, 2017).

During the evolution of therapeutic resistance, both bacteria and cancer can exhibit a phenomenon termed collateral sensitivity, where resistance to one drug aligns with sensitivity to another drug (Pluchino et al., 2012; Pál et al., 2015; Hall et al., 2009). Likewise, collateral resistance occurs when resistance to one drug aligns with resistance to another drug. The relationship between genotype (e.g. gene expression, somatic mutations, etc.) and fitness of a cell line can be represented by a fitness landscape. In the case of drug response, we define fitness as the EC50 of a cell line to a given drug, where increasing EC50 denotes higher fitness in the presence of this drug. EC50 is defined as the concentration of drug which achieves half-maximal effective response. Of importance, a cell line with the same genotype may have varying fitnesses (EC50s) under the selection pressure of different drugs.

In collateral resistance, the fitness landscapes of the organism (bacteria or cancer) in the presence of each drug would show “positive correlation” (Nichol et al., 2019). This is because genotypic changes that cause increased fitness in presence of the first drug also allow increased fitness in the presence of the second drug as well. Next, comparing fitness landscapes in the setting of collateral sensitivity will show “negative correlation,” where genotypic changes leading to increased fitness in the presence of the first drug will cause decreased fitness in the presence of the second drug. Finally, comparing fitness landscapes in the presence of different treatments will not always demonstrate clear positive or negative correlation. Instead, the evolution of resistance to one drug may lead to variable changes in response to the second drug. In this setting, the evolutionary landscapes would be “uncorrelated.” Here, predictive models would be especially useful in treatment planning, as relative collateral sensitivity or resistance cannot be inferred based solely on treatment history.

In the case of collateral sensitivity, a clinician could ideally control disease progression by switching to a collaterally sensitive drug whenever resistance develops. Even if the illness was never completely eradicated, the pathogen or neoplasm would be dampened enough to minimize harm to the patient. Yet evolution is rarely so easy to predict. Several studies have aimed to identify examples of collateral sensitivity in either bacteria or cancer, and many have shown that exposure to identical therapies has resulted in different responses between evolutionary replicates (Imamovic and Sommer, 2013; Munck et al., 2014; Nichol et al., 2015; Zhao et al., 2016; Dhawan et al., 2017; Nichol et al., 2019; Maltas and Wood, 2019). Additionally, Zhao et al., examined changes in collateral sensitivity in acute lymphoblastic leukemia (ALL) over time (Zhao et al., 2016). Here, they produced temporal collateral sensitivity maps to show how drug response evolved over time and between evolutionary replicates (Zhao et al., 2016). Although Zhao et al. did examine these changes through time, many collateral sensitivity experiments compare only initial and final drug response after resistance to the primary treatment has evolved (Dhawan et al., 2017; Nichol et al., 2019). These intermediate steps are crucial for determining whether the evolution of therapeutic resistance leads to a collateral fitness landscape that is consistently positively/negatively correlated or uncorrelated through time.

For the first time in a solid tumor, we examine the repeatability of collateral sensitivity across time as cells evolve resistance to standard treatment. In doing so, we use two EWS cell lines, A673 and TTC466. The A673 cell line contains the *t(11;22)* translocation resulting in the *EWSR1/FLI1* gene fusion (Szuhai et al., 2006; Martinez-Ramirez et al., 2003). This fusion is the most common genetic aberration found in 90-95% EWS tumors (Sankar et al., 2014; Szuhai et al., 2006). In contrast, the TTC466 cell line has a *t(21;22)* translocation resulting in the EWS-ERG gene fusion, which only occurs only in 5-10% of EWS tumors (Sankar et al., 2014; Szuhai et al., 2006). After splitting the cell lines into evolutionary replicates, they were exposed to standard chemotherapy and their response to a panel of drugs was assessed over time. All drugs included in this study may be found in Table 1. We hypothesize that Ewing’s sarcoma cell lines repeatedly exposed to standard chemotherapy will demonstrate divergent evolutionary paths between evolutionary replicates despite nearly identical experimental conditions and initial genotype. Finding patterns of collateral resistance (positively correlated landscapes), sensitivity (negatively correlated landscapes) or variation (uncorrelated landscapes) within these divergent paths may provide useful insight in exploring new treatment options for EWS patients.

**Table 1:** All drugs referenced in the study, their abbreviations, and classifications.

| Drug Name | Abbreviation | Class |
| --- | --- | --- |
| Dactinomycin | ActD | Antineoplastic antibiotic |
| Cyclophosphamide | Cyclo | Alkylating agent |
| Ifosfamide | Ifo | Alkylating agent |
| Doxorubicin | Doxo | Anthracycline |
| Etoposide | Etp | Topoisomerase II inhibitor |
| Olaparib | Ola | PARP inhibitor |
| Pazopanib | Paz | Tyrosine kinase inhibitor |
| Vorinostat | SAHA | Histone deacetylase inhibitor |
| Irinotecan<br>(active metabolite) | SN38 | Topoisomerase I inhibitor |
| SP-2509 | SP | Lysine-specific demethylase<br>1 inhibitor |
| Temozolomide | TMZ | Alkylating agent |
| Vincristine | Vin | Alkaloid |
| Sodium thiosulfate | NaThio | Drug activation reagent |

## Results

### The long term evolution of therapeutic resistance

This work examines the evolution of collateral sensitivity and resistance in two EWS cell lines during repeated exposure to a standard chemotherapy regimen over time. At the onset of the experiment, each cell line was split into eight evolutionary replicates, five experimental and three control. Due to contamination, Replicate 2 from the A673 cell line was excluded from the analysis, leaving four experimental and three control replicates in this cell line. Each experimental evolutionary replicate then underwent the same drug cycling, as demonstrated in Figure 1. Briefly, experimental replicates were incubated in cycles of vincristine-doxorubicin-cyclophosphamide (VDC) and etoposide-cyclophosphamide (EC) combinations (Grier et al., 2003). This procedure models standard-of-care given to EWS patients, which consists of cycles of vincristine-doxorubucin-cyclophosphamide (VDC) and etoposide-ifosfamide (EI) combinations. Because ifosfamide requires metabolic activation and no activated compound is commercially available, we chose to substitute ifosfamide with cyclophosphamide, as these compounds are analogs (Fleming, 1997). Control replicates were maintained in only vehicle control. More details can be found in Transparent Methods.

**Figure 1:**
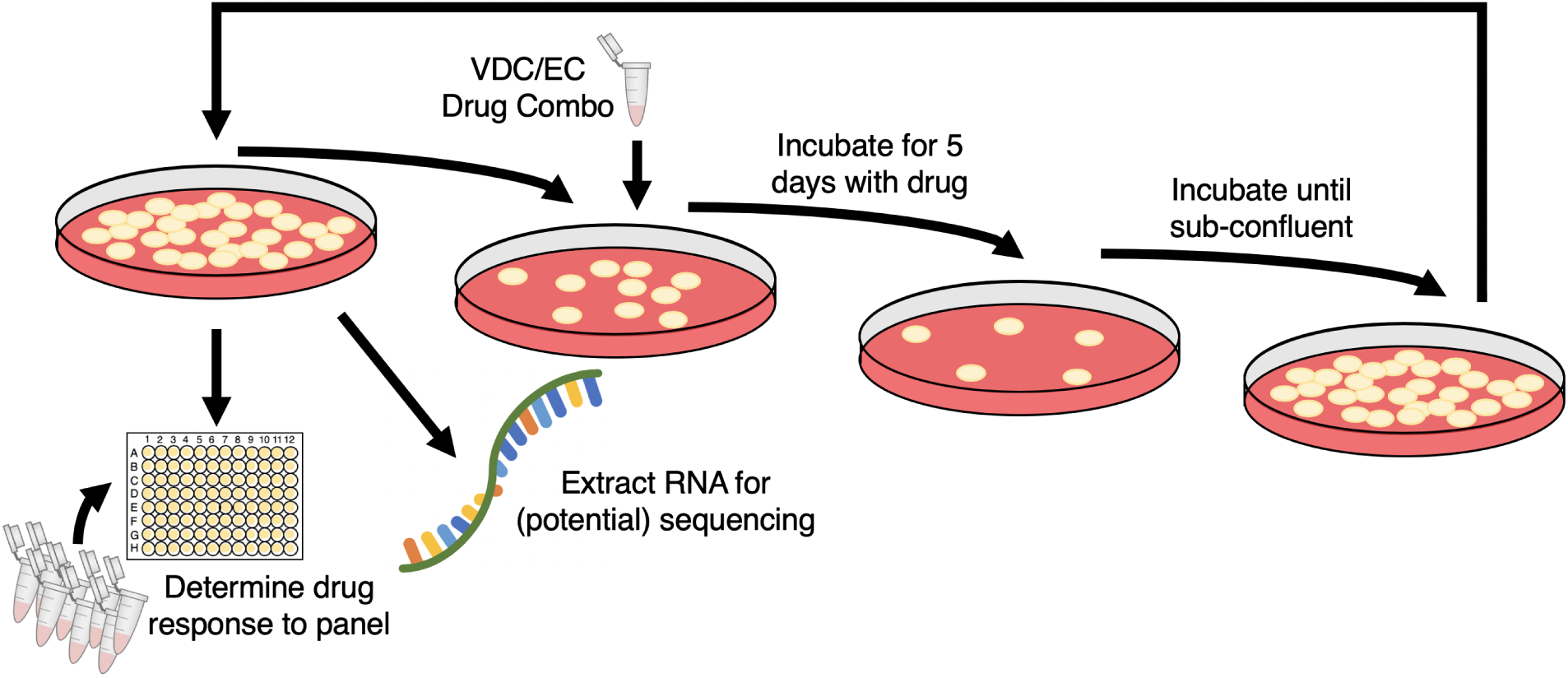
Overview of experimental evolution of resistance in Ewings sarcoma cell lines. As cells recovered from each exposure, cells were tested for their sensitivity for a panel of drugs and samples were frozen for potential use in RNA-sequencing. The drug dosage was only increased once throughout the experiment, at the fifth exposure to the VDC combination, described in Transparent Methods. Additionally, drug toxicity assays are performed at each time point to evaluate changes in therapeutic resistance or sensitivity over time. Although each cell line began with 5 experimental and 3 control evolutionary replicates, the A673 cell line lost one experimental replicate due to contamination.

After proliferating to sub-confluent density in maintenance medium, a fraction of cells from all evolutionary replicates were snap frozen for RNA extraction, another fraction underwent drug sensitivity assays to 12 drugs, and another fraction of the cells were plated for exposure to the next cycle of the alternate drug combination. For each drug at each time point, the EC50 of each evolutionary replicate was derived by fitting the drug-response triplicate data to a four-parameter log-logistic model as described in Transparent Methods. A plot of all dose-response triplicates with their estimated EC50 can be found in the linked GitHub repository.

### Discerning changes in chemo-sensitivity and -resistance across time

Figures 2 and 3 display the changes in drug response to 9 agents over time in the A673 cell line. In addition to the nine drugs displayed this figure, two additional drugs (Pazopanib and Vincristine) and a drug activation reagent (sodium thiosulfate) were included in the drug sensitivity assays. Data for these drugs are included in Supplemental Figures 1 and 2.

**Figure 2:**
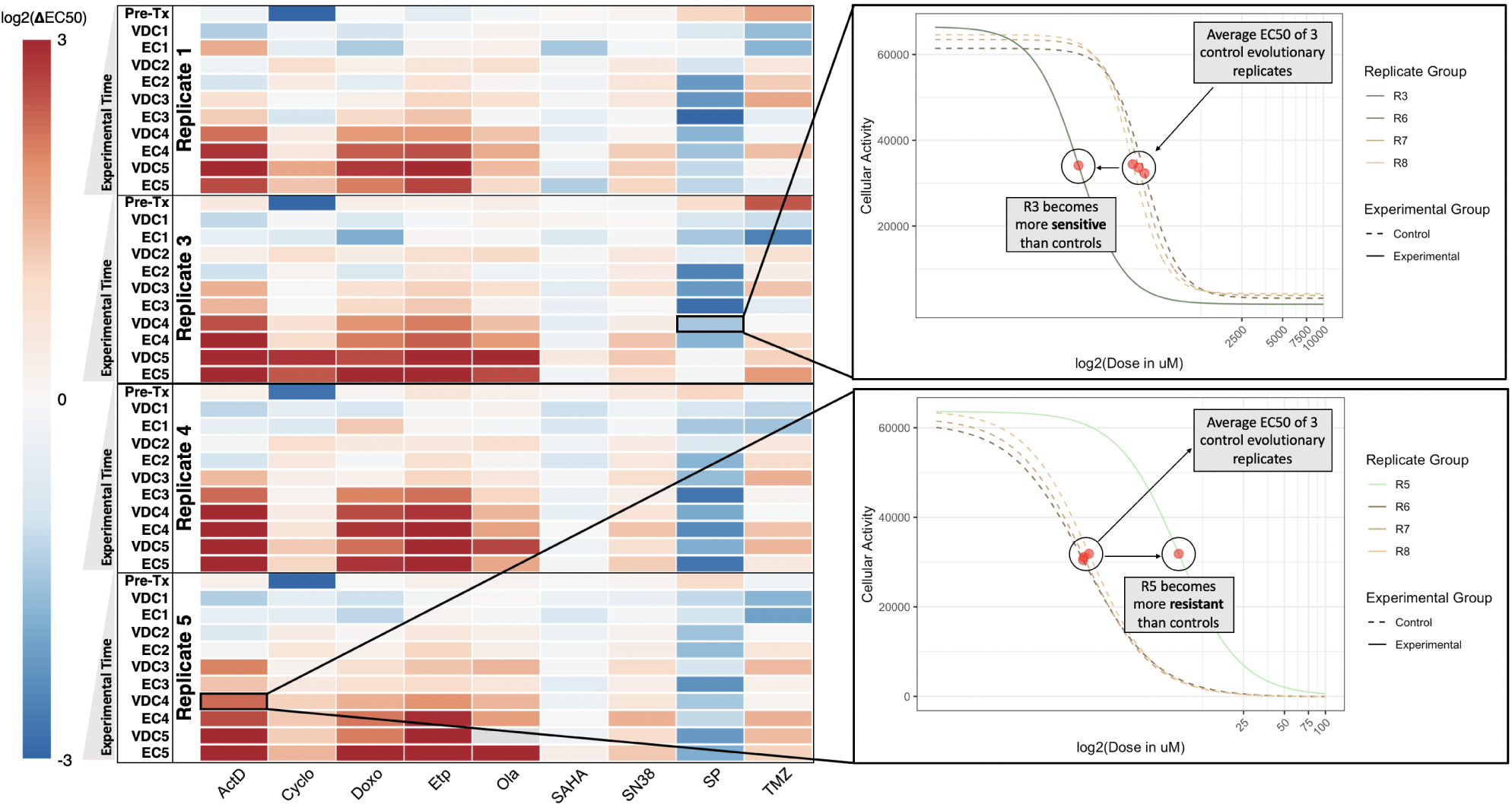
Temporal collateral sensitivity map representing EC50 changes to a panel of drugs as the A673 cell line develops resistance to standard treatment. **Left:** A heatmap representing how the EC50 to a panel of nine drugs changes in 4 A673 cell line evolutionary replicates as they are exposed to the VDC/EC drug combinations over time. Color represents the log_2_ fold change of EC50 to a drug (columns) for a replicate at a given evolutionary time point (rows) compared to the average EC50 of the three control evolutionary replicates at the corresponding time point. Values above log_2_(3) or below log_2_(−3) are represented by log_2_(3) and log_2_(−3), respectively. Time points are denoted as the drug combination that a given replicate has recently recovered from. For example, the data representing dose-response models after the first application of the VDC drug combination would be labeled with VDC1. Of note, the EC50 of olaparib in Replicate 5 at the VDC5 timepoint is indeterminate due to a poorly fit dose-response model. This value in the heatmap is denoted as gray, but Supplementary Figure 1 remains uncensored. **Right:** Top, a plot of the dose-response curves for Replicate 3 and all control replicates (Replicates 6, 7, 8) in response to SP-2509 (SP) at the VDC4 time point. Bottom, a plot of the dose-response curve for Replicate 5 and all control replicates in response to dactinomycin at the VDC4 time point. Cellular activity is measured by enzymatic conversion of alamarBlue, normalized to background florescence. Estimated EC50 for each replicate is denoted with a red circle. These two dose-response plots demonstrate how the heatmap (left) values were calculated, where the control EC50 values are averaged and the heatmap values represent the log_2_ fold change between a given replicate and this mean EC50 value.

**Figure 3:**
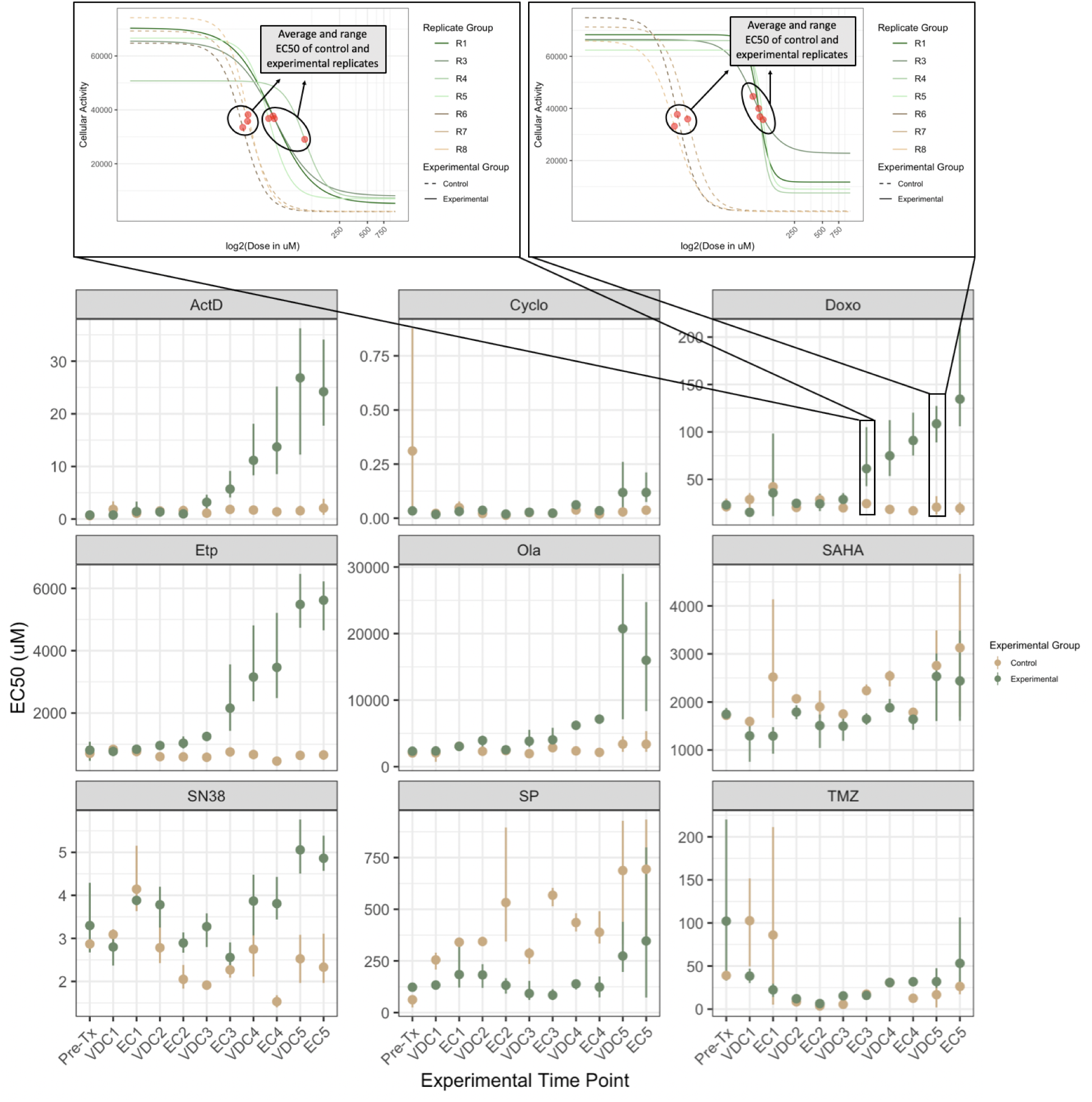
Point-range plots demonstrating EC50 changes in A673 experimental and control replicates over time. **Bottom:** Point-range plots representing the changes in drug response to a panel of nine drugs. Experimental time points (x-axis) represent which step in the drug cycle the replicates have just recovered from. Points on the plot represent the average EC50 for the group, either experimental or control. Lines represent the range for the entire group. The EC50 of olaparib for Replicate 5 after the fifth exposure to VDC is indeterminate due to a poorly fit dose-response model, and has been removed from this drug’s VDC5 time point experimental group EC50 average and range calculations. This value has not been censored in Supplemental Figures 1 and 2. The y-axis of all the point-range plots has uM units, except Cyclo, where the unit is percent of chemically activated 4-hydroxycyclophosphamide solution by volume. **Top:** Two plots demonstrating a more detailed view of the dose-response data represented at the EC3 and VDC5 time points in the Doxo point-range chart. Cellular activity is measured by enzymatic conversion of alamarBlue, normalized to background florescence. Comparing these two plots shows the clear divergence in drug response between experimental and control evolutionary replicates as the treatment regimen continued.

Unlike the A673 cell line, the TTC466 develops resistance to only a few of the tested drugs and when it is developed, it is not maintained across evolutionary time. It is unsurprising, therefore, to see that Supplementary Figures Supplemental Figures 3 and 4 demonstrate that changes in collateral sensitivity responses are not significantly different between experimental and control replicates for the TTC466 cell line. As such, we focus our analysis on the A673 cell line, and the heatmap and point-range plots for the TTC466 cell line can be found in Supplemental Figures 3 and 4, respectively.

Figure 2 shows a temporal collateral sensitivity map which represents the log_2_ fold change of EC50 to a drug (columns) for an experimental replicate at a given time point (rows) compared to the average EC50 of the three control evolutionary replicates to the same drug, at that time point. In the top right of the figure, we see the drug response of Replicate 3 to SP-2509 after its fourth exposure to VDC (VDC4), along with the three control evolutionary replicates at this time point. This example demonstrates a move towards sensitivity in the experimental replicate. In the bottom right, we can interrogate Replicate 5 at the same time point in response to dactinomycin, where the EC50 of this experimental replicate is more resistant than the control replicates. The temporal collateral sensitivity maps found in Supplementary Figures 1 and 3 include the log_2_ fold change for each control evolutionary replicate from the average of the three control evolutionary replicates at the corresponding time point. Ideally, this value will be close to zero (white), because the three control replicates should have similar EC50s.

Figure 3 contains point-range plots for the nine drugs included in our analysis demonstrating the average and range of EC50 values for experimental and control replicates at each time point in the experiment. In response to doxorubicin, the control replicates remained stable across each progressive time point, but the experimental replicates became increasingly more resistant as they were repeatedly exposed to standard treatment. Examining these point-range plots also allows us to observe the overall stability of drug response in control replicates, which are not being evolved under the selection pressure of the VDC/EC drug cycling. For example, the EC50 range for control replicates to dactinomycin is so minimal that the lines representing range are not visible for most time points in the ActD panel of Figure 3. On the other hand, control replicates show significant variation in their response to SN38 and temozolomide (TMZ) at many time points.

The point-range plots in Figure 3 also allow for a more nuanced interrogation of the log-ratio changes displayed in Figure 2. For example, although all experimental replicates appear to become more sensitive in response to SP in Figure 2, the point-range plots in Figure 2 demonstrate that the control evolutionary replicates are becoming more resistant over time, while the experimental evolutionary replicates remain somewhat stable. This relative sensitivity between experimental and control evolutionary replicates is visualized as a negative log_2_ fold change in Figure 2. Yet, because the control, not experimental, evolutionary replicates changed over time, it is less clear whether or not this negative log-ratio should be interpreted as a move towards sensitivity in the experimental replicates. Early in the treatment series, the experimental and control replicates have very similar average EC50 measurements, demonstrating divergence over time. It is likely that the control conditions caused increased resistance to SP in the cell lines (leading to increasing EC50 in control replicates), while the application of the VDC/EC drug cycling regimen promoted sensitivity (leading to the appearance of stable EC50 through time).

### Surveying the stochasticity of evolution

While examining Figures 2 and 3, we see predictable development of collateral resistance to some drugs, but evolutionary stochasticity and divergence in collateral response between replicates was observed in the response to others. For example, the cells were initially sensitive to dactinomycin and moved into a distinct state of collateral resistance in all replicates. This leads us to the preliminary conclusion that the evolutionary landscape of the cells under the VDC/EC selection pressure and the landscape of the cells under the dactinomycin selection pressure would show strong positive correlation. Next, all replicates acquired relatively consistent resistance to doxorubucin and etoposide over time, which is to be expected, because these two reagents are included in the treatment regimen. All replicates relatively consistently evolved from sensitive to resistant in response to olaparib, SN38, and temozolomide. In response to vorinostat (SAHA), all replicates appear to show minor sensitivity across time, but no discernible trend in toward greater sensitivity nor resistance. Finally, there was moderate sensitivity to SP seen in all A673 replicates, again with no discernible trends through time. Due to the variation in response to SAHA or SP over time, we would conclude that the fitness landscapes of cells exposed to these drugs compared to the landscapes of cells exposed to VDC/EC are relatively uncorrelated.

Unexpectedly, most replicates acquired only mild resistance to cyclophosphamide, a drug which is included in both cycles of the treatment regimen. Additionally, one of the control evolutionary replicates has an outlier EC50 estimate, which causes the pre-treatment (Pre-Tx) experimental evolutionary replicates to appear more sensitive to cyclophosphamide than the control evolutionary replicates. Although this outlier replicate is likely experimental noise, as it does not remain an outlier at future time points, the dose-response model is well fit to the data and we determined that censoring this replicate’s time point EC50 estimate would be inappropriate.

### Differential gene expression analysis provides insight into the mechanisms of drug response

Eighteen samples from the A673 cell line were RNA-sequenced, visualized in the left panel of Figure 4. All the samples were ranked based on their response to the 12 drugs included in the drug sensitivity panels. These rankings are visually represented in waterfall plots of the log_2_ fold change in EC50 for all sequenced samples against all drugs can be seen in Supplementary Figures 5-16. For each drug, differential gene expression (DE) analysis was performed between samples that rank in the top and bottom third of responses towards the drug. Results for each drug’s DE analyses, including the analyses highlighted below, may be found in Supplementary Information.

**Figure 4:**
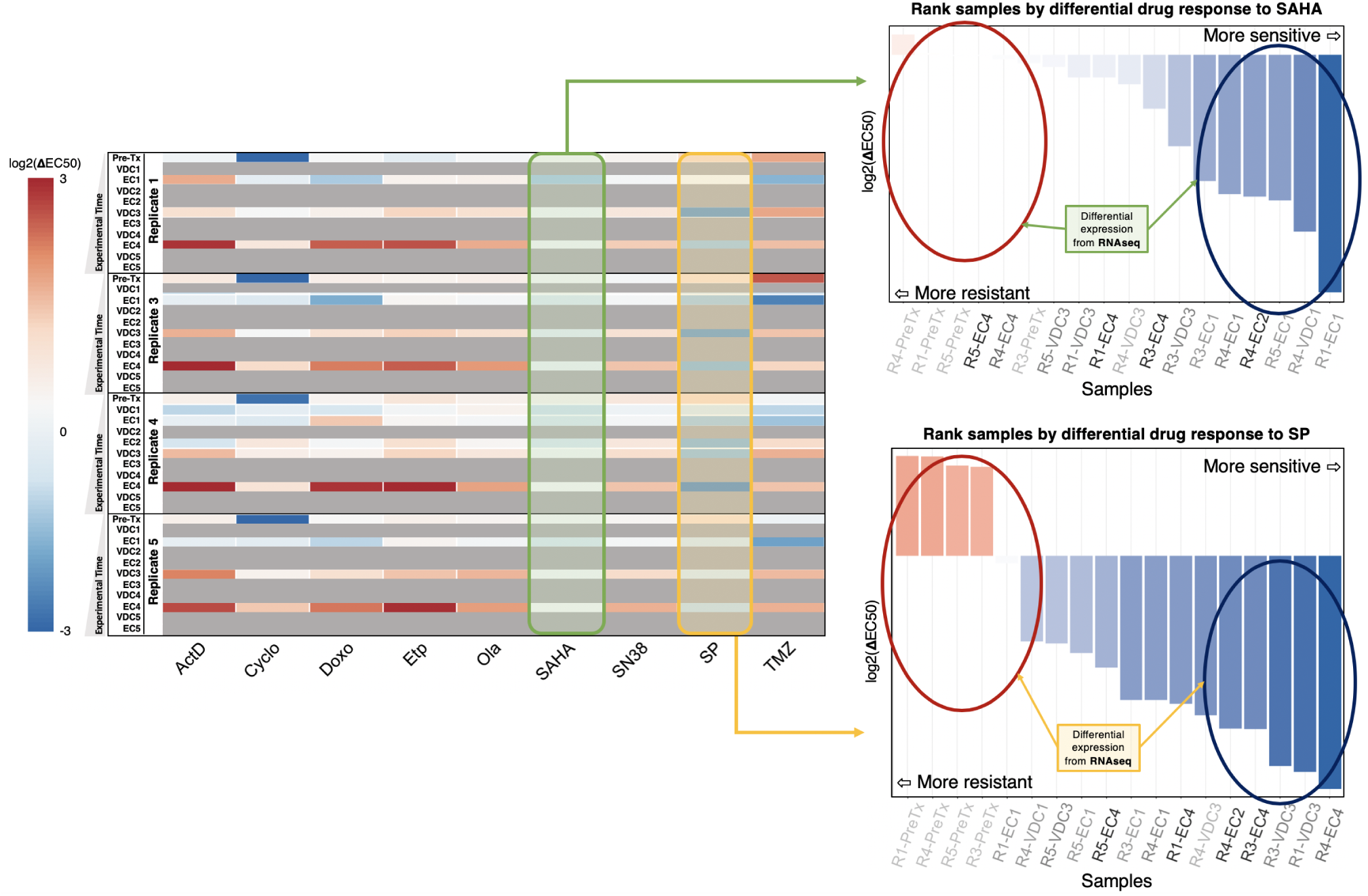
RNA-sequencing and differential gene expression analysis provide insight into states of collateral sensitivity and resistance. **Left:** The temporal collateral sensitivity map from Figure 2, where all samples that were not sequenced are overlayed with gray. **Right:** Two waterfall plots representing the samples ranked by their responses to the two drugs, vorinostat (SAHA, top) and SP-2509 (SP, bottom). Sample labels on the x-axis are represented by darker colors the longer they have been evolved in the evolutionary experiment.

In many cases, it is clear that ranking samples by their change in drug response also ranks them based on how long they’ve been exposed to the treatment regimen. Although this is not unexpected, interpreting the DE results in this context becomes more difficult. Significant differences in gene expression may be related to a sample’s chemosensitivity/chemoresistance, but causation cannot be inferred, because these differentially expressed genes may simply be altered in response to continued exposure to the treatment regimen. We chose to highlight the DE analyses where ranking samples in response to a given drug did not consistently arrange them in the order that they were isolated from the drug-cycling treatment. To that end, the waterfall plots in Supplementary Figures 5-16 have darker sample labels (x-axis) depending on how long they’ve been exposed to the treatment regimen (e.g. a sample label from the IE2 time point will be lighter than a sample from the IE3 time point). This makes it easier to visualize whether the time points are well distributed in the log_2_ fold change rankings.

The right panel of Figure 4 demonstrates how samples were ranked based on their response to vorinostat (SAHA) and SP-2509 (SP). Genes with significantly increased expression in a SAHA-resistant or SAHA-sensitive state are listed in Table 2, while genes with significantly increased expression in an SP-resistant or SP-sensitive state are listed in Table 3.

**Table 2:** Genes with significant differential expression between SAHA-resistant and SAHA-sensitive samples. Differential gene expression analysis was performed using EBSeq in R, with maxround set to 15 and FDR of 0.05.

| Genes with ↑ expression in<br>SAHA-sensitive state | | | |
| --- | --- | --- | --- |
| ACOT | ACPP | AHR | B3GNT5 |
| CCL2 | FOS | GAL | NUP188 |
| RN7SL5P | SCNN1G | TRAV5 |  |
| Genes with ↑ expression in<br>SAHA-resistant state | | | |
| ABCB1 | KAZALD1 | RPS26 | SMAD6 |
| TRGC1 |  |  |  |

**Table 3:** Genes with significant differential expression between SP-resistant and SP-sensitive samples. Differential gene expression analysis was performed using EBSeq in R, with maxround set to 15 and FDR of 0.05.

| Genes with ↑ expression in SP-sensitive state | | | |
| --- | --- | --- | --- |
| ALX1 | AMZ2 | APOBEC3C | ARHGEF6 |
| CD63 | DCN | FAM72D | FAM92A |
| HIST1H1T | IL33 | IRX3 | LINC00326 |
| LITAF | LYN | MRPS18C | NPIPA5 |
| NRG1 | PCSK6 | PTGR1 | PYCARD |
| RTN | SP100 | SSTR1 | TMEM192 |
| TSPAN5 | YAF2 | ZFAND1 | ZNF277 |
| Genes with ↑ expression in SP-resistant state | | | |
| ADGRL2 | ANKS6 | AP3B2 | AP5Z1 |
| ARHGEF9 | C7 | CAMKV | CCAR2 |
| CD24 | CDH4 | CHGA | CORO7 |
| CRMP1 | DGCR8 | DHCR7 | DPP3 |
| EPHA4 | FASN | FOXO3B | FRG2FP |
| GALNS | HBA2 | HDAC10 | INCENP |
| INTS1 | KSR1 | LIN28B | LINC01089 |
| LRCH2 | MAN2C1 | MEG3 | MEG8 |
| MRGPRF | MRNIP | MSRA | NEB |
| NEFM | NOM1 | NUP210 | PBX1 |
| PC | PCBP2-OT1 | PCDH17 | PLXNB1 |
| PPP1R1B | PRRC2B | PTPRG-AS1 | PPYGO1 |
| RNF130 | RNF44 | SBNO2 | SCAMP4 |
| SCARA3 | SLC16A7 | SLC29A2 | SLITRK3 |
| SYK | TAF15 | TAF1C | TAF6L |
| TMEM271 | TUBB3 | VAX1 | WDR17 |
| WDR27 | ZNF354C | ZNF414 | ZNF667 |
| ZNF667-AS1 | ZNF675 | ZNF730 | ZNF736 |

## Discussion

In this work, we evolved two EWS cell lines, A673 and TTC466, with repeated exposure to standard-of-care chemotherapy in order to investigate the evolution of collateral sensitivity and resistance through time. We produced a temporal collateral sensitivity map to examine the drug sensitivity assays for nine of these drugs through time in the A673 cell line (see Figure 2). Likewise, Figure 3 demonstrates how the average and range EC50 between A673 experimental replicates and control replicates diverged as the experimental replicates continued the drug cycling treatment regimen. Supplementary Figures 1 and 2 contain the drug response changes to all 12 drugs, with no censored data. Supplementary Figures 3 and 4 also exhibit the changes in drug response within the TTC466 cell line; however, the main text focuses on the A673 cell line due to minimal shifts in resistance observed in response to the treatment regimen in the TTC466 cell line. Figure 2 shows that as the A673 experimental replicates were repeatedly exposed to the treatment regimen, states of collateral resistance emerge consistently towards some drugs, while responses to other agents remain variable. For example, despite no exposure to the drug, all replicates consistently moved to a state of collateral resistance towards dactinomycin, providing an example of positively correlated evolutionary landscapes. On the other hand, all replicates demonstrate possible collateral sensitivity to SP over time, although these results are not conclusive as the control evolutionary replicates moved towards resistance more than the experimental evolutionary replicates moved towards sensitivity. All replicates show variable minimal collateral sensitivity to SAHA, but no state of collateral resistance or sensitivity dominates for many time points or between replicates. Both of these drugs would have uncorrelated evolutionary landscapes in comparison to the landscape under the VDC/EC selection pressure. Additionally, these results imply that although there are consistent changes that allow for collateral resistance to dactinomycin, these changes do not invariably cause a consistent pattern of collateral sensitivity to SAHA or SP. When collateral sensitivity or resistance cannot be consistently identified, gene signatures or other predictive models would be especially helpful in treatment planning.

We also note that the A673 experimental replicates consistently become more resistant to doxorubucin and etoposide, and there is a moderate shift towards resistance to cyclophosphamide. All of these drugs, in addition to vincristine, are included in the drug combination treatment regimen, which makes the evolution of resistance unsurprising. Although response to the drug combinations (VDC and EC) themselves was not measured, increased resistance to the treatment regimen was demonstrated by the decreased time required to reach sub-confluence after each cycle as time. As seen in Methods, this necessitated an increase in the final VDC drug combination concentration. Importantly, drug sensitivity changes in response to drug combinations may be the result of a variety of changes in response to the individual drug combination components, and this analysis would have been improved by measuring response to the drug combination triplicates and duplicates through time.

After analyzing the repeatability (or lack thereof) of the evolution of collateral sensitivity and resistance in the A673 EWS cell line, 18 samples from across various time points were sent for RNA-sequencing (Figure 4). We identified significantly increased expression of ABCB1 (also known as MDR1) in the state of SAHA-resistance, seen in Table 2. This gene has previously been implicated in chemotherapeutic multidrug resistance (Chen and Sikic, 2012). Additionally, CCL2 was found to have increased expression in a SAHA-sensitive state. Using an *in vitro* experiment, Gatti and Sevko et al. describe how adding SAHA to the temozolomide treatment of melanoma may stymie cancer growth by interfering with CCL2 signaling. This is consistent with increased CCL2 expression leading to SAHA sensitivity, as cells that are more reliant on CCL2 signaling could experience a stronger effect from its disruption (Gatti et al., 2014).

Next, we see a greater number of differentially expressed genes when examining response to SP than SAHA. SP inhibits lysine-specific demethylase 1 (LSD1, also known as KDMA1), which primarily acts as a histone demethylase (O’Leary et al., 2016). Increased expression of LSD1 has been implicated in many types of cancers (e.g. breast, prostate), and its targeted inhibition is being investigated for therapeutic potential in EWS (Yang et al., 2018). Due to the novelty of LSD1 inhibitors (including SP-2509), there is very little known regarding genomic biomarkers of sensitivity or resistance. In Table 3, however, we see some notable trends in the significantly differentially expressed genes between good and poor responders to SP. First, many zinc protein fingers, which often play a role in transcriptional regulation, have increased expression in both SP-sensitive and -resistant states (Klug, 1999). Additionally, three TATA-box-binding-protein (TBP) associated factor (TAF) proteins have increased expression in SP-resistant states. Again, these genes are implicated in transcriptional regulation (Hampsey and Reinberg, 1997). Although these results do not imply any one mechanism for SP-sensitivity or -resistance, it is evident that the regulation of gene expression may play a significant role in the response to this drug.

As noted previously, states of collateral sensitivity and resistance are often not immutable. Instead, these states are frequently the result of many fleeting evolutionary contingencies. For instance, Nichol et al. demonstrated that after *E. coli* evolved resistance to cefoxatime (a *β*-lactam antibiotic) in 60 evolutionary replicates, there were highly heterogeneous changes in collateral sensitivity and resistance to alternative antibiotics (Nichol et al., 2019). Genotypic heterogeneity was discovered as well, with five variants of the *β*-lactamase gene which likely played a role in the variable drug responses. Additionally, Dhawan et al. derived cell lines of ALK-positive non-small cell lung cancer, where each cell line was resistant to a secondline therapy (Dhawan et al., 2017). Subsequently, the cell lines were exposed to the same panel of second-line treatments in an effort to identify drug combinations that elicit collateral sensitivity. The study found that collateral sensitivity was most often evolved towards etoposide and pemetrexed. Although these drugs had the most optimal response, it was inconsistent, leading to the conclusion that collateral sensitivity is a dynamic state, which is a ‘moving target’ instead of a predictable outcome.

With this understanding, our experiment would, of course, benefit from even more experimental evolutionary replicates to confirm the repeatability of some observations. For example, the evolution of collateral resistance to dactinomycin in all four A673 experimental replicates is consistently stable in the data presented here. However, given the vast genetic contingencies that lead to changes in drug response, observing said stability over many additional replicates would provide a more convincing argument for the consistent evolution of dactinomycin collateral resistance following exposure to VDC/EC.

Performing this experiment in a greater number of cell lines would provide improved insight into the spectrum of responses across various EWS cases. Using increased replicates and additional cell lines would also enhance the analysis of differential gene expression results between responders and non-responders, as we could examine whether differentially expressed genes are consistently dysregulated across different cell lines after exposure to the same experimental conditions. Finally, this work could be improved by examining how collateral drug response in EWS changes during relaxed selection after many drug cycles of the treatment regimen have been applied (Card et al., 2019). This would represent a model that is even more consistent with refractory EWS in a clinical setting, as patients will often have a gap between the initial standard treatment and the selection of second-line treatments.

Despite these caveats, this work provides valuable insight into the evolution of collateral resistance and sensitivity in EWS throughout exposure to standard treatment. Although, many studies have examined the role that collateral sensitivity and resistance play in therapeutic response, they frequently ignore intermediate time points during the development of resistance to a primary treatment. In this work, we aimed to examine collateral sensitivity and resistance across time during development of therapeutic resistance to EWS standard-of-care. We believe this is the first temporal map of collateral sensitivity and resistance in a solid tumor cell line. Using this map, we can see that the path towards collateral sensitivity or resistance is not always straight. Likewise, evolutionary replicates can demonstrate varying collateral drug responses through time, even if their drug response near the end of the experiment is similar. Gene expression signatures can provide clarity when choosing a new treatment in the setting of a tumultuous trajectory towards the evolution of collateral sensitivity or resistance.

## Limitations of the Study

Our experiment would benefit from even more experimental evolutionary replicates to confirm the repeatability of some observations. For example, the evolution of collateral resistance to dactinomycin in all four A673 experimental replicates is consistently stable in the data presented here. However, given the vast genetic contingencies that lead to changes in drug response, observing said stability over many additional replicates would provide a more convincing argument for the consistent evolution of dactinomycin collateral resistance following exposure to VDC/EC. Performing this experiment in a greater number of cell lines would provide improved insight into the spectrum of responses across various EWS cases. Using increased replicates and additional cell lines would also enhance the analysis of differential gene expression results between responders and non-responders, as we could examine whether differentially expressed genes are consistently dysregulated across different cell lines after exposure to the same experimental conditions. Finally, this work could be improved by examining how collateral drug response in EWS changes during relaxed selection after many drug cycles of the treatment regimen have been applied (Card et al., 2019). This would represent a model that is even more consistent with refractory EWS in a clinical setting, as patients will often have a gap between the initial standard treatment and the selection of second-line treatments.

## Resource Availability

### Lead Contact

Jacob Scott is the lead contact for this manuscript. He can be reached at.

### Materials Availability

We are glad to share A673 cell line and its resistant derivatives generated in this study with a completed MTA.

### Data and Code Availability

Gene expression data may be accessed from the Sequence Read Archive with the BioProject accession number: PRJNA630293. Drug response data from the long term evolution experiments may be accessed through Mendeley Data DOI: 10.17632/vbc3gjjgyn.1. The code to perform all analyses is available via GitHub at https://github.com/jessicascarborough/ES-CS-evolution.

## Acknowledgements

JGS would like to thank the NIH Loan Repayment Program for their generous support, the Paul Calabresi Career Development Award for Clinical Oncology (NIH K12CA076917), the Carson Sarcoma Foundation, and Chemowarrior Foundation. Additionally, JAS thanks the NIH for support through the T32GM007250 grant. The authors thank the Lerner Research Institute Genomic Core for RNA-sequencing and Kathleen Pishas for helpful discussions and for sharing the cell lines.

## Author contributions statement

JAS performed data processing, wrote all associated code, analyzed the data, and wrote the manuscript. EM and MH performed the long term evolution experiments, drug sensitivity assays, RNA extraction, and wrote the manuscript. A. Dhawan contributed to experimental design and analyzed the data. A. Durmaz performed RNA-sequencing quality control and alignment. PA contributed to experimental design and analyzed the data. JGS analyzed the data and wrote the manuscript. These contributions are graphically illustrated in Figure 5.

**Figure 5:**
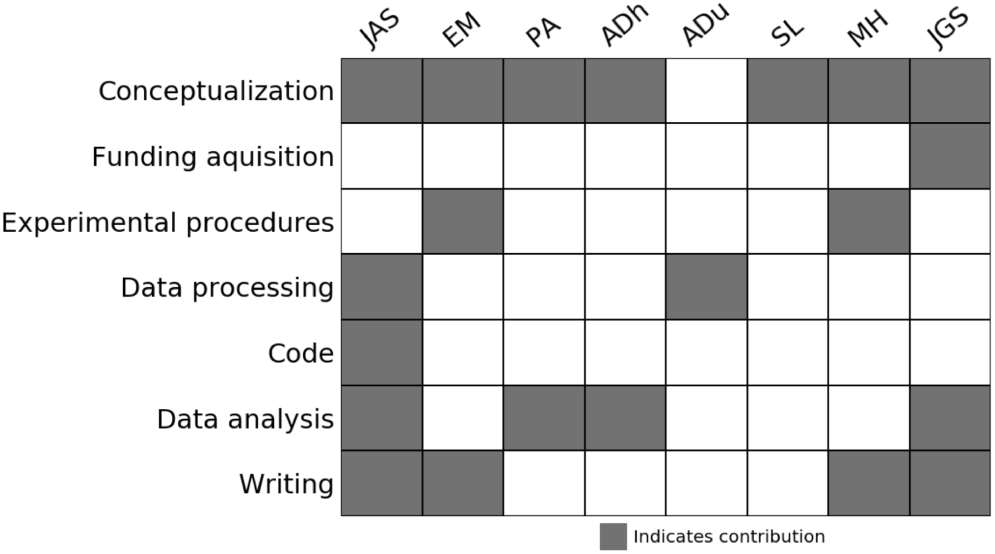
Author Contributions.

## References

Ahmed, A.A., Zia, H., Wagner, L., 2014. Therapy resistance mechanisms in ewing’s sarcoma family tumors. Cancer chemotherapy and pharmacology 73, 657–663.

Card, K.J., LaBar, T., Gomez, J.B., Lenski, R.E., 2019. Historical contingency in the evolution of antibiotic resistance after decades of relaxed selection. PLoS biology 17.

Chen, K.G., Sikic, B.I., 2012. Molecular pathways: regulation and therapeutic implications of multidrug resistance. Clinical cancer research 18, 1863–1869.

Dhawan, A., Nichol, D., Kinose, F., Abazeed, M.E., Marusyk, A., Haura, E.B., Scott, J.G., 2017. Collateral sensitivity networks reveal evolutionary instability and novel treatment strategies in alk mutated non-small cell lung cancer. Scientific Reports 7, 1–9.

Esiashvili, N., Goodman, M., Marcus, R.B., 2008. Changes in incidence and survival of ewing sarcoma patients over the past 3 decades: Surveillance epidemiology and end results data. Journal of pediatric hematology/oncology 30, 425–430.

Fleming, R.A., 1997. An overview of cyclophosphamide and ifosfamide pharmacology. Pharmacotherapy: The Journal of Human Pharmacology and Drug Therapy 17, 146S–154S.

Gatti, L., Sevko, A., De Cesare, M., Arrighetti, N., Manenti, G., Ciusani, E., Verderio, P., Ciniselli, C.M., Cominetti, D., Carenini, N., et al., 2014. Histone deacetylase inhibitor-temozolomide co-treatment inhibits melanoma growth through suppression of chemokine (cc motif) ligand 2-driven signals. Oncotarget 5, 4516.

Grier, H.E., Krailo, M.D., Tarbell, N.J., Link, M.P., Fryer, C.J., Pritchard, D.J., Gebhardt, M.C., Dickman, P.S., Perlman, E.J., Meyers, P.A., et al., 2003. Addition of ifosfamide and etoposide to standard chemotherapy for ewing’s sarcoma and primitive neuroectodermal tumor of bone. New England Journal of Medicine 348, 694–701.

Hall, M.D., Handley, M.D., Gottesman, M.M., 2009. Is resistance useless? multidrug resistance and collateral sensitivity. Trends in pharmacological sciences 30, 546–556.

Hampsey, M., Reinberg, D., 1997. Transcription: why are tafs essential? Current Biology 7, R44–R46.

Huang, M., Lucas, K., 2010. Current therapeutic approaches in metastatic and recurrent ewing sarcoma. Sarcoma 2011.

Hunold, A., Weddeling, N., Paulussen, M., Ranft, A., Liebscher, C., Jürgens, H., 2006. Topotecan and cyclophosphamide in patients with refractory or relapsed ewing tumors. Pediatric blood & cancer 47, 795–800.

Imamovic, L., Sommer, M.O., 2013. Use of collateral sensitivity networks to design drug cycling protocols that avoid resistance development. Science translational medicine 5, 204ra132–204ra132.

Klug, A., 1999. Zinc finger peptides for the regulation of gene expression. Journal of molecular biology 293, 215–218.

Maltas, J., Wood, K.B., 2019. Pervasive and diverse collateral sensitivity profiles inform optimal strategies to limit antibiotic resistance. PLoS biology 17.

Martinez-Ramirez, A., Rodriguez-Perales, S., Meléndez, B., Martinez-Delgado, B., Urioste, M., Cigudosa, J., Benitez, J., 2003. Characterization of the a673 cell line (ewing tumor) by molecular cytogenetic techniques. Cancer genetics and cytogenetics 141, 138–142.

Munck, C., Gumpert, H.K., Wallin, A.I.N., Wang, H.H., Sommer, M.O., 2014. Prediction of resistance development against drug combinations by collateral responses to component drugs. Science translational medicine 6, 262ra156–262ra156.

Nichol, D., Jeavons, P., Fletcher, A.G., Bonomo, R.A., Maini, P.K., Paul, J.L., Gatenby, R.A., Anderson, A.R., Scott, J.G., 2015. Steering evolution with sequential therapy to prevent the emergence of bacterial antibiotic resistance. PLoS computational biology 11.

Nichol, D., Rutter, J., Bryant, C., Hujer, A.M., Lek, S., Adams, M.D., Jeavons, P., Anderson, A.R., Bonomo, R.A., Scott, J.G., 2019. Antibiotic collateral sensitivity is contingent on the repeatability of evolution. Nature communications 10, 1–10.

O’Leary, N.A., Wright, M.W., Brister, J.R., Ciufo, S., Haddad, D., McVeigh, R., Rajput, B., Robbertse, B., Smith-White, B., Ako-Adjei, D., et al., 2016. Reference sequence (refseq) database at ncbi: current status, taxonomic expansion, and functional annotation. Nucleic acids research 44, D733–D745.

Pál, C., Papp, B., Lázár, V., 2015. Collateral sensitivity of antibiotic-resistant microbes. Trends in microbiology 23, 401–407.

Pluchino, K.M., Hall, M.D., Goldsborough, A.S., Callaghan, R., Gottesman, M.M., 2012. Collateral sensitivity as a strategy against cancer multidrug resistance. Drug Resistance Updates 15, 98–105.

Ries, L.A.G., 1999. Cancer incidence and survival among children and adolescents: United States SEER program, 1975-1995. 99, National Cancer Institute.

Sankar, S., Theisen, E.R., Bearss, J., Mulvihill, T., Hoffman, L.M., Sorna, V., Beckerle, M.C., Sharma, S., Lessnick, S.L., 2014. Reversible lsd1 inhibition interferes with global ews/ets transcriptional activity and impedes ewing sarcoma tumor growth. Clinical cancer research 20, 4584–4597.

Scott, J., Marusyk, A., 2017. Somatic clonal evolution: A selection-centric perspective. Biochimica et Biophysica Acta (BBA)-Reviews on Cancer 1867, 139–150.

Szuhai, K., Ijszenga, M., Tanke, H.J., Rosenberg, C., Hogendoorn, P.C., 2006. Molecular cytogenetic characterization of four previously established and two newly established ewing sarcoma cell lines. Cancer genetics and cytogenetics 166, 173–179.

Womer, R.B., West, D.C., Krailo, M.D., Dickman, P.S., Pawel, B.R., Grier, H.E., Marcus, K., Sailer, S., Healey, J.H., Dormans, J.P., et al., 2012. Randomized controlled trial of interval-compressed chemotherapy for the treatment of localized ewing sarcoma: a report from the children’s oncology group. Journal of Clinical Oncology 30, 4148.

Yang, G.J., Lei, P.M., Wong, S.Y., Ma, D.L., Leung, C.H., 2018. Pharmacological inhibition of lsd1 for cancer treatment. Molecules 23, 3194.

Zhao, B., Sedlak, J.C., Srinivas, R., Creixell, P., Pritchard, J.R., Tidor, B., Lauffenburger, D.A., Hemann, M.T., 2016. Exploiting temporal collateral sensitivity in tumor clonal evolution. Cell 165, 234–246.

