## Supplementary Information for "Identifying states of collateral sensitivity during the evolution of therapeutic resistance in Ewing’s sarcoma"

### 1. Supplementary Figures and Legends

#### *1.1. Complete A673 EC50 Data*

The following plots mirror Figures 2 and 3, respectively. The data does not censor the EC50 for Replicate 5 against olaparib at the VDC5 timepoint, as seen in Figures 2 and 3. Additionally, the drugs removed from main text analysis, vincristine, pazopanib, and sodium thiosulfate are included. These drugs were censored in the main text due to poorly fit dose-response models. Interpretation of these plots can be found in the main text.

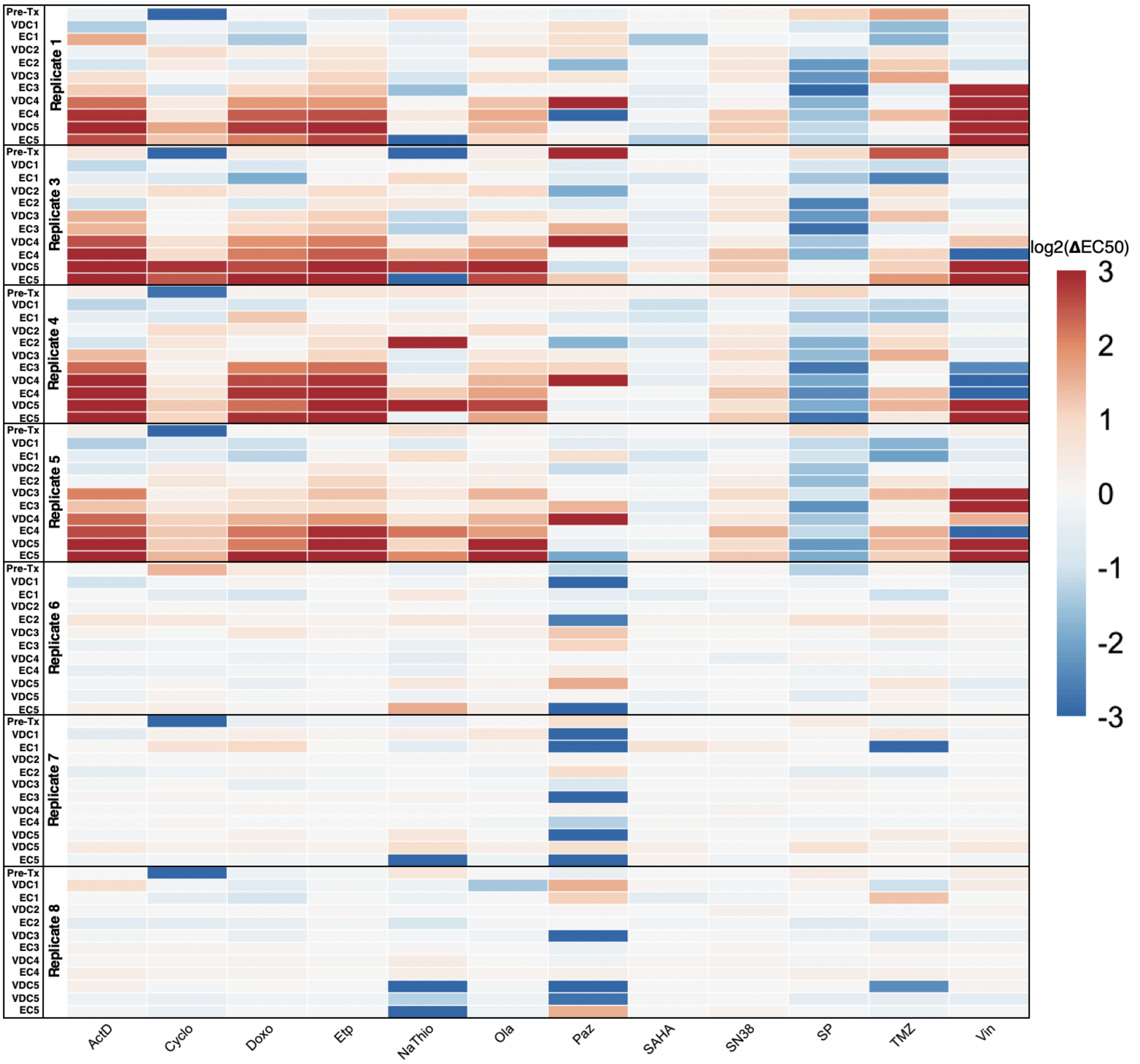


Supplementary Figure 1: **Uncensored temporal collateral sensitivity map representing EC50 changes to panel of drugs in A673 cell line as it develops resistance to standard treatment, related to Figure 2.** A heatmap representing how the EC50 to a panel of nine drugs changes in 4 experimental and 3 control evolutionary replicates from the A673 cell line as they are exposed to the VDC/EC drug combinations over time. Color represents the $\log_2$ fold change of EC50 to a drug (columns) for a replicate at a given evolutionary time point (rows) compared to the average EC50 of the three control evolutionary replicates at the corresponding time point. Time points are denoted as the drug combination that a given replicate has recently recovered from. For example, the data representing dose-response models after the first application of the VDC drug combination would be labeled with VDC1.

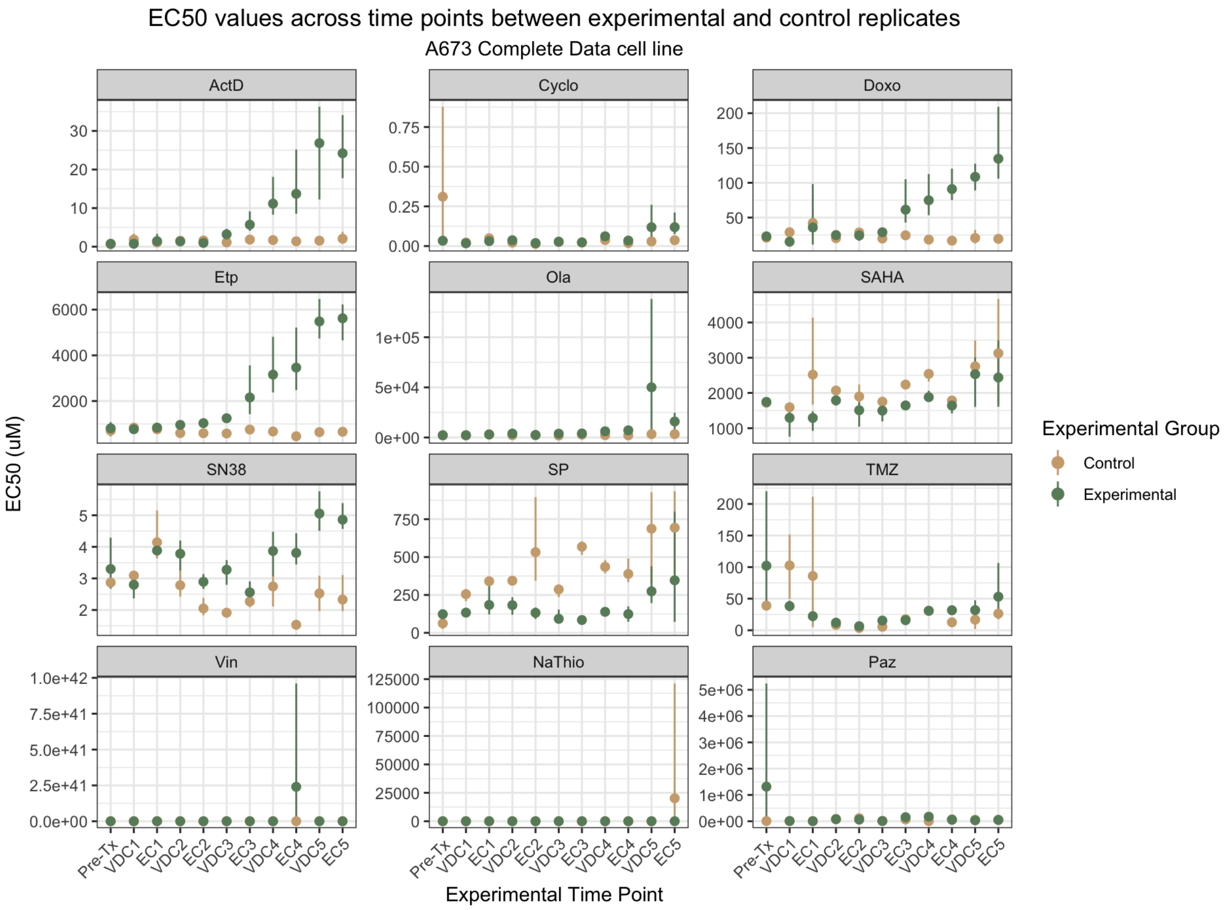


Supplementary Figure 2: **Uncensored point-range plots demonstrating EC50 changes in A673 experimental and control replicates over time, related to Figure 3.** Point-range plots representing the changes in drug response to a panel of 12 drugs. Experimental time points (x-axis) represent which step in the drug cycle the replicates have just recovered from. Points on the plot represent the average EC50 for the group, either experimental or control. Lines represent the range for the entire group. The y-axis of all the point-range plots has uM units, except Cyclo and NaThio, where the units are percent by volume.

#### 1.2. *Complete TTC466 EC50 Data*

Supplementary Figures 3 and 4 displays the changes in drug response to all 12 agents over time in the TTC466 cell line. In comparing the two cell lines, it is clear that the A673 cell line displays more stable behavior, while TTC466 shows much more variability over time. In other words, as the treatment cycles progress, the A673 cell line tends to move steadily towards a resistant or sensitive state, while the TTC466 cell line tends to fluctuate more. The TTC466 control replicates also tend to have more fluctuation between time points, despite being exposed to only media and having relative agreement between technical replicates. For this reason, we chose to focus our analysis on the A673 cell line, while the TTC466 cell line results can be found in the Supplementary Information.

In Supplementary Figure 3, we see that after the first exposure to the VDC drug combination (VDC1) in the TTC466 cell line, resistance to cyclophosphamide suddenly emerges. This doesn't occur in any other replicate, nor at any other time point. These findings were confirmed by examining the drug-response curve at this time point to ensure a well-fit model. Two hypotheses for why the replicate didn't retain the cyclophosphamide-resistant trait in the next generation include an equally rapid loss of this trait in the next generations or a bottleneck selection during the procedure where the cells that were resistant to cyclophosphamide were not plated for the next round of the drug treatment cycle. Next, another example of drug-response fluctuation in the TTC466 cell line may have been mistaken as a rare shift in drug response if only one evolutionary replicate had been performed. In Supplementary Figure 3, we see that after the second exposure to the EC combination (EC2), the EC50 of every experimental replicate has increased chemoresistance to olaparib before returning to a more sensitive state after the next drug cycle. Supplementary Figure 4, demonstrates that there is a large range in the control replicates at the corresponding time point, which makes the comparison between the experimental and control replicates less reliable; however, it is clear that from the time points before and after EC2, the EC50 increases significantly at EC2.

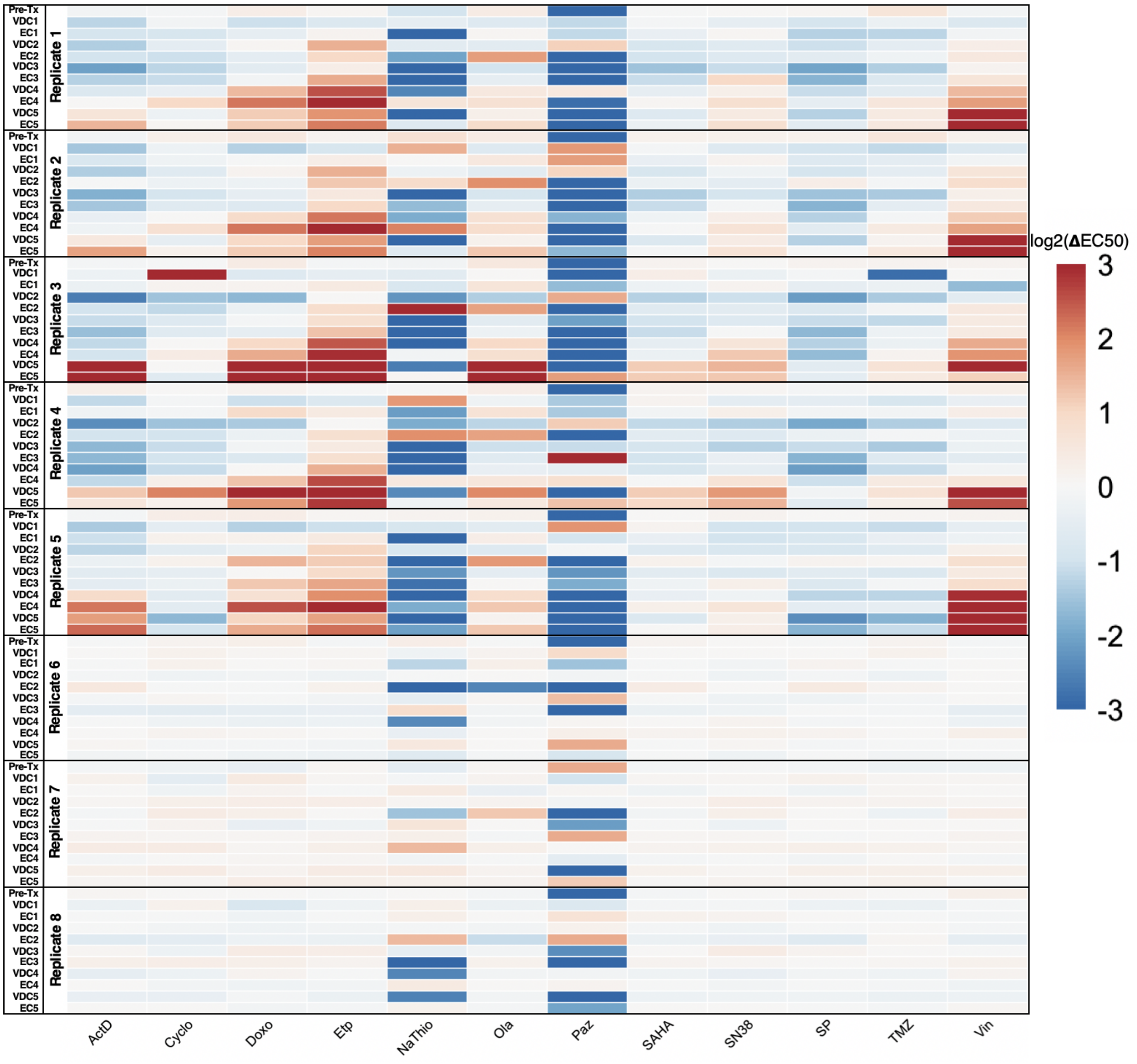


Supplementary Figure 3: **Uncensored temporal collateral sensitivity map representing EC50 changes to panel of drugs in TTC466 cell line as it develops resistance to standard treatment, related to Figure 2.** A heatmap representing how the EC50 to a panel of nine drugs changes in 5 experimental and 3 control evolutionary replicates from the TTC466 cell line as they are exposed to the VDC/EC drug combinations over time. Color represents the $\log_2$ fold change of EC50 to a drug (columns) for a replicate at a given evolutionary time point (rows) compared to the average EC50 of the three control evolutionary replicates at the corresponding time point. Time points are denoted as the drug combination that a given replicate has recently recovered from. For example, the data representing dose-response models after the first application of the VDC drug combination would be labeled with VDC1.

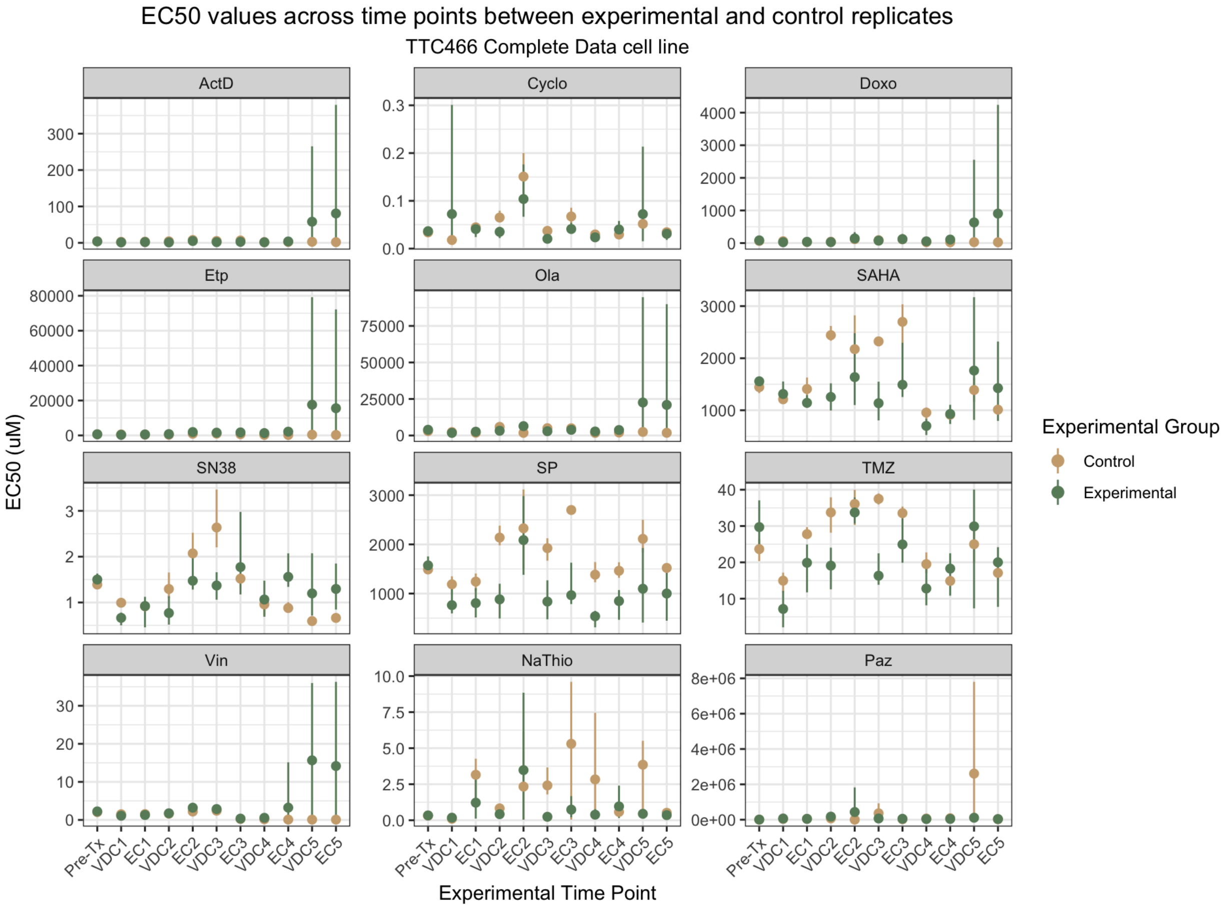


Supplementary Figure 4: **Uncensored point-range plots demonstrating EC50 changes in TTC466 experimental and control replicates over time, related to Figure 3.** Point-range plots representing the changes in drug response to a panel of 12 drugs. Experimental time points (x-axis) represent which step in the drug cycle the replicates have just recovered from. Points on the plot represent the average EC50 for the group, either experimental or control. Lines represent the range for the entire group. The y-axis of all the point-range plots has uM units, except Cyclo and NaThio, where the units are percent by volume.

*1.3. Waterfall plots for sequenced samples against all drugs*

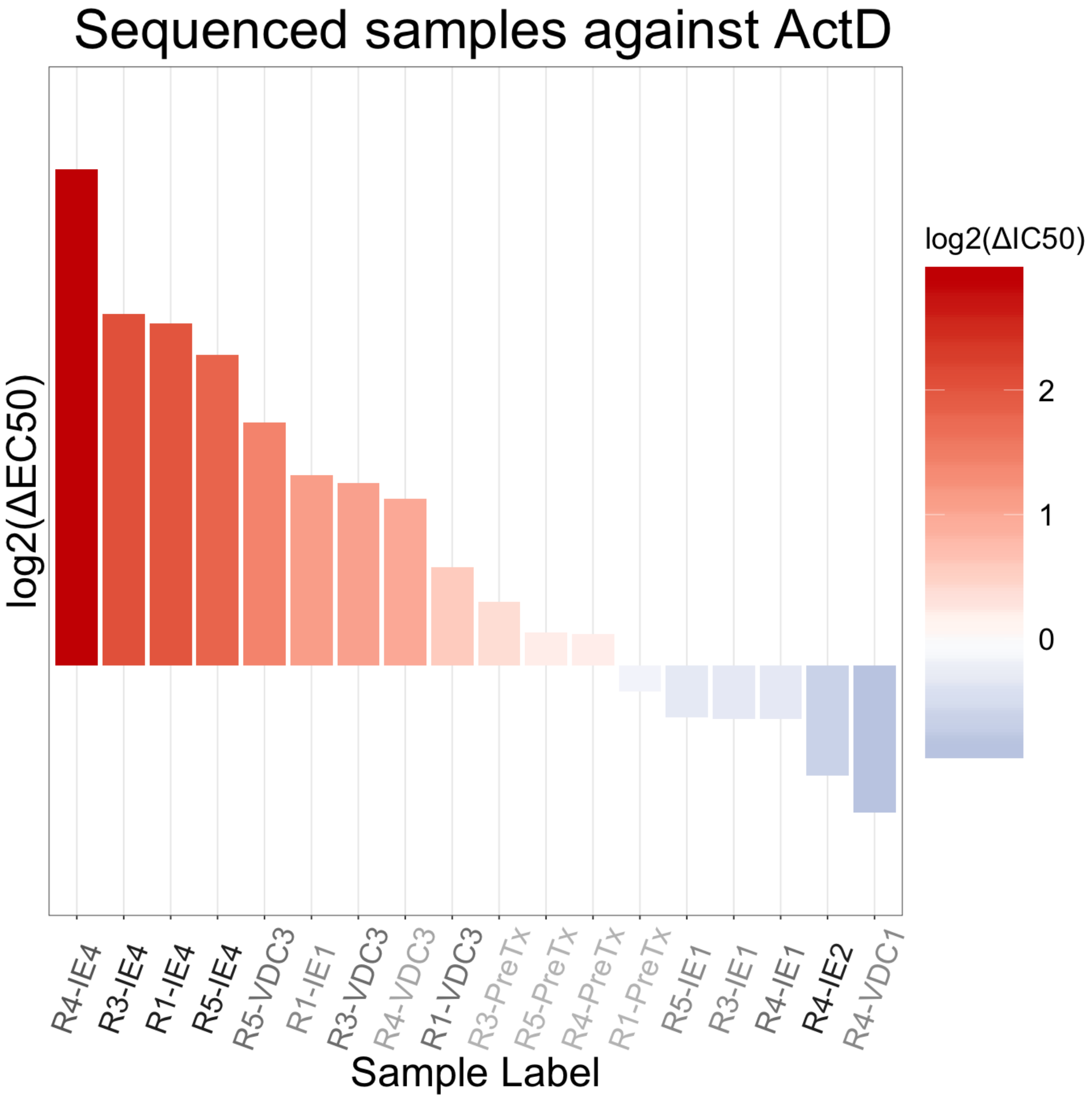


Supplementary Figure 5: **Waterfall of EC50 values for sequenced samples against dactinomycin, related to Figure 4.** Color represents $\log_2$ change in EC50 between the sample and average control EC50 at the given time point. Red shows a change towards resistance, while blue shows a change towards sensitivity. Samples are ranked along the x-axis from least-to-most sensitive. Sample labels on the x-axis are represented by darker colors the longer they have been evolved in the evolutionary experiment.

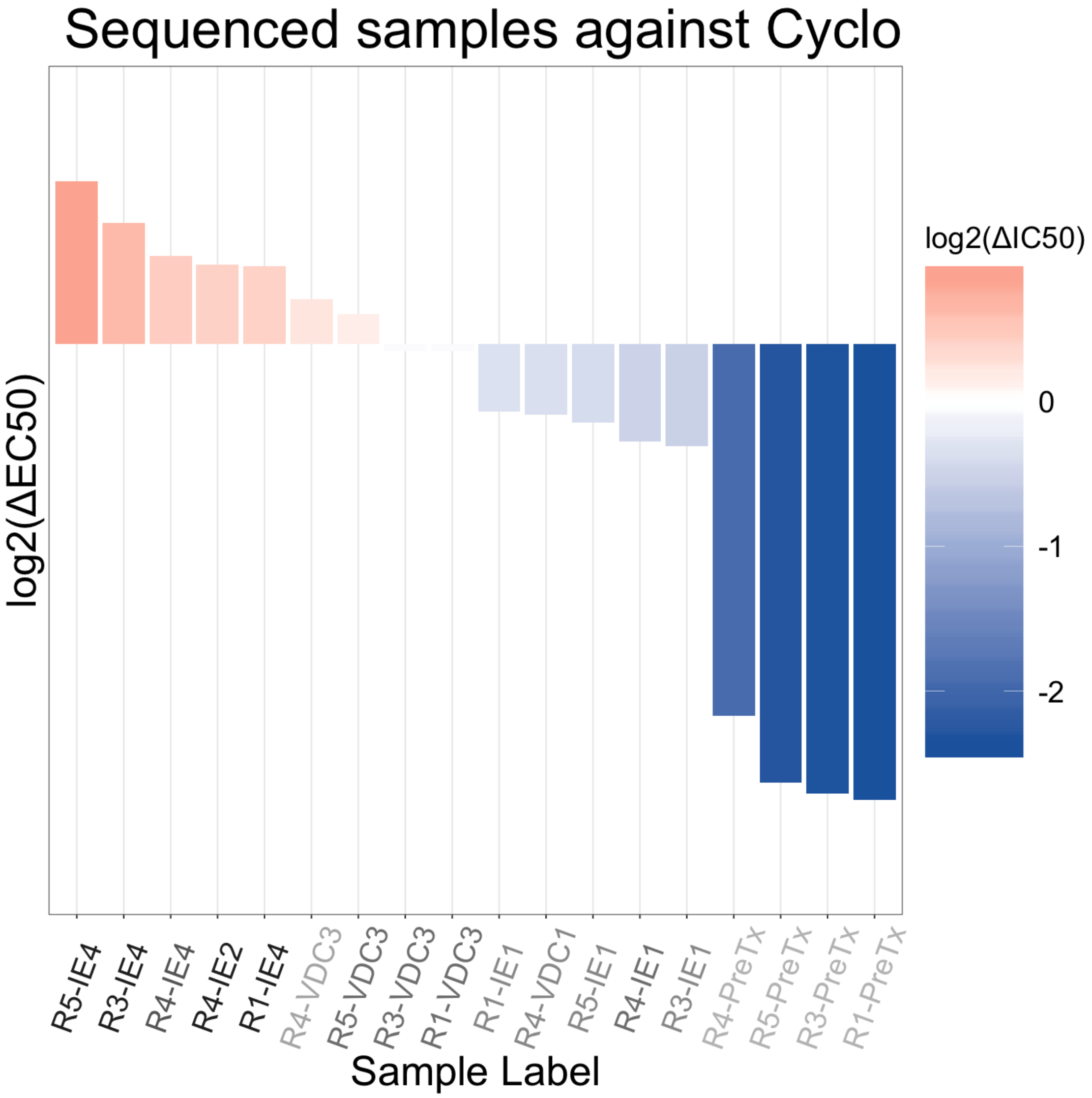


Supplementary Figure 6: **Waterfall of EC50 values for sequenced samples against cyclophosphamide, related to Figure 4.** Color represents $\log_2$ change in EC50 between the sample and average control EC50 at the given time point. Red shows a change towards resistance, while blue shows a change towards sensitivity. Samples are ranked along the x-axis from least-to-most sensitive. Sample labels on the x-axis are represented by darker colors the longer they have been evolved in the evolutionary experiment.

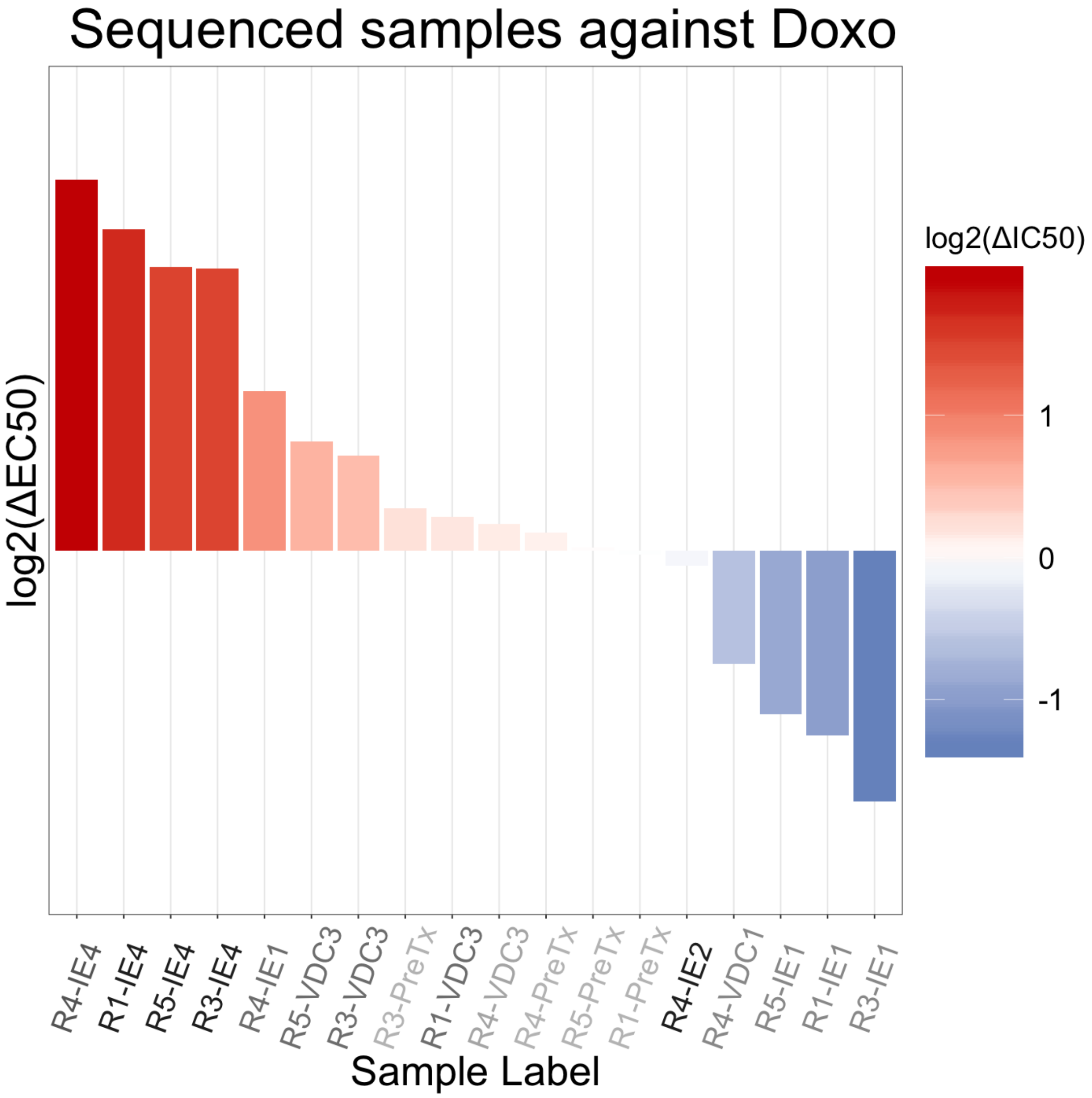


Supplementary Figure 7: **Waterfall of EC50 values for sequenced samples against doxorubicin, related to Figure 4.** Color represents $\log_2$ change in EC50 between the sample and average control EC50 at the given time point. Red shows a change towards resistance, while blue shows a change towards sensitivity. Samples are ranked along the x-axis from least-to-most sensitive. Sample labels on the x-axis are represented by darker colors the longer they have been evolved in the evolutionary experiment.

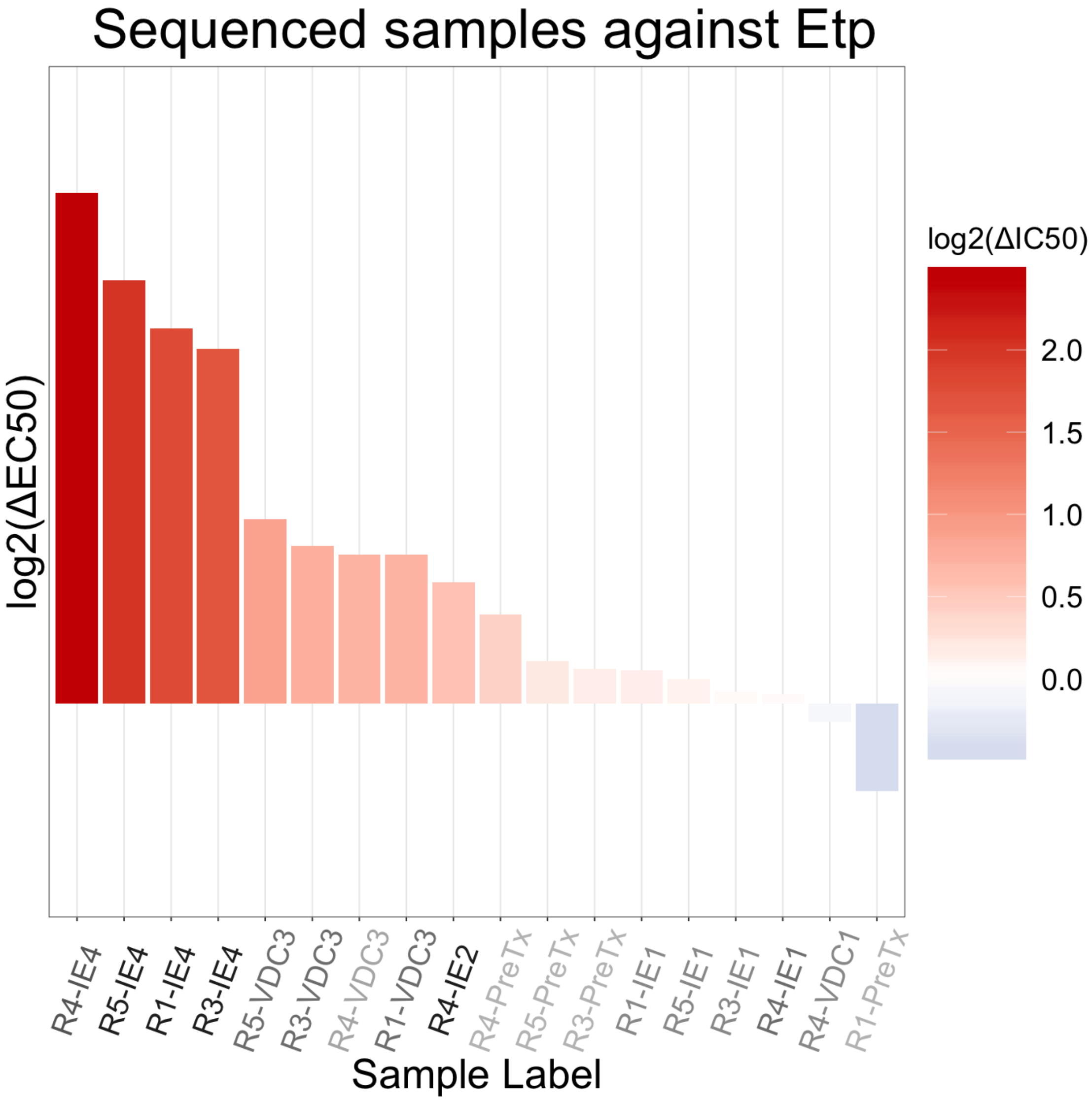


Supplementary Figure 8: **Waterfall of EC50 values for sequenced samples against etoposide, related to Figure 4.** Color represents $\log_2$ change in EC50 between the sample and average control EC50 at the given time point. Red shows a change towards resistance, while blue shows a change towards sensitivity. Samples are ranked along the x-axis from least-to-most sensitive. Sample labels on the x-axis are represented by darker colors the longer they have been evolved in the evolutionary experiment.

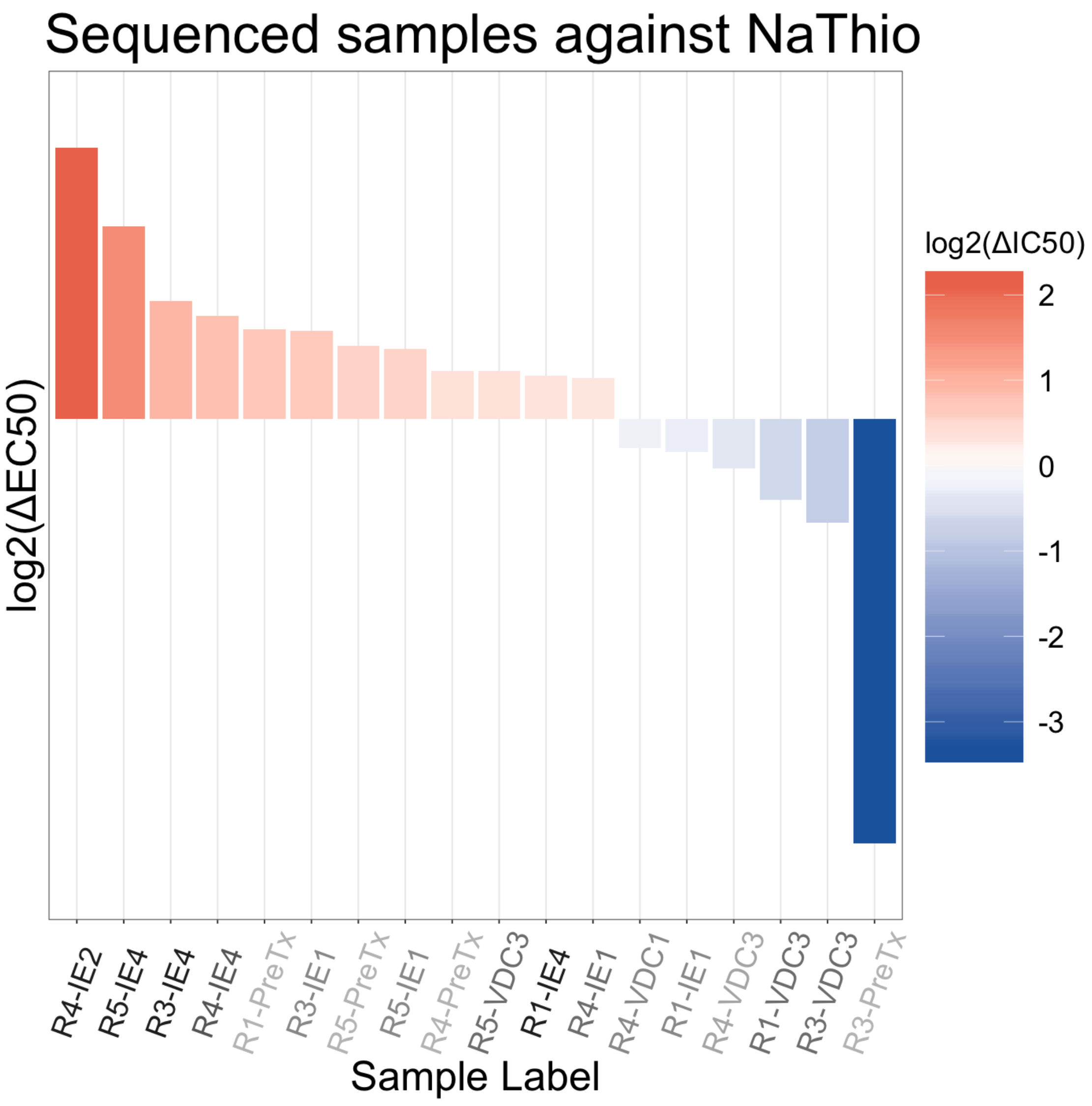


Supplementary Figure 9: **Waterfall of EC50 values for sequenced samples against sodium thiosulfate, related to Figure 4.** Color represents $\log_2$ change in EC50 between the sample and average control EC50 at the given time point. Red shows a change towards resistance, while blue shows a change towards sensitivity. Samples are ranked along the x-axis from least-to-most sensitive. Sample labels on the x-axis are represented by darker colors the longer they have been evolved in the evolutionary experiment.

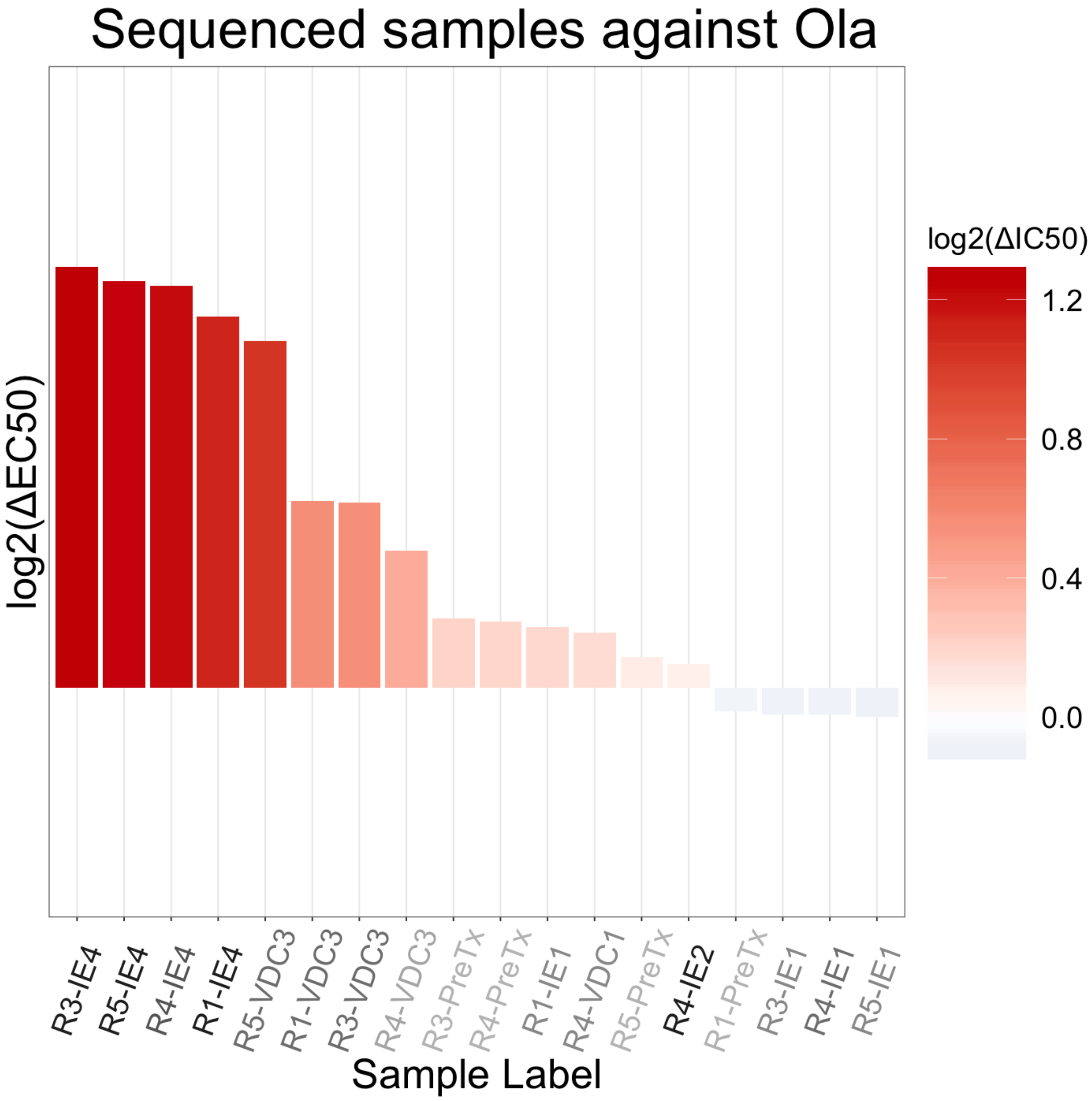


Supplementary Figure 10: **Waterfall of EC50 values for sequenced samples against olaparib, related to Figure 4.** Color represents $\log_2$ change in EC50 between the sample and average control EC50 at the given time point. Red shows a change towards resistance, while blue shows a change towards sensitivity. Samples are ranked along the x-axis from least-to-most sensitive. Sample labels on the x-axis are represented by darker colors the longer they have been evolved in the evolutionary experiment.

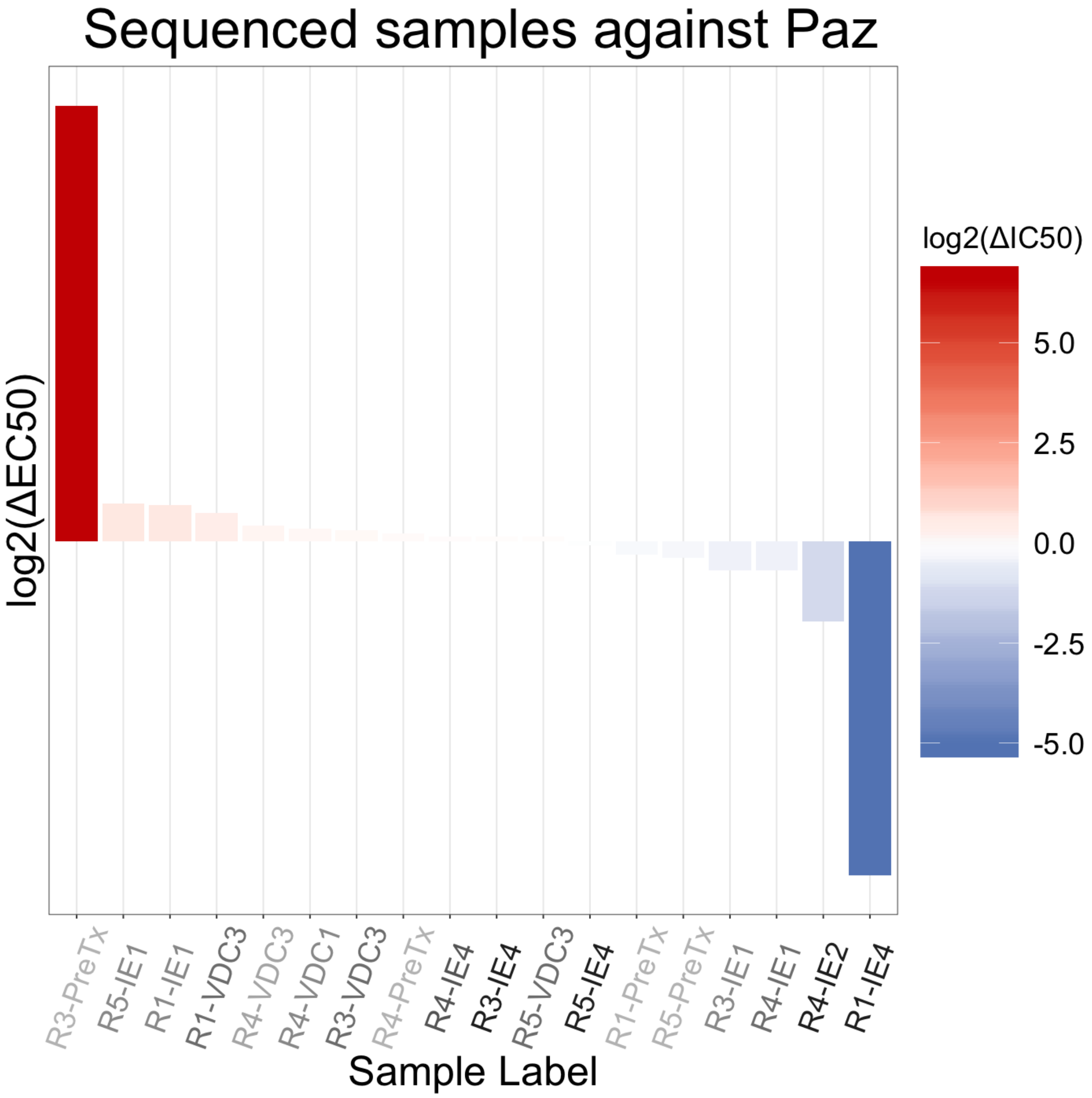


Supplementary Figure 11: **Waterfall of EC50 values for sequenced samples against pazopanib, related to Figure 4.** Color represents $\log_2$ change in EC50 between the sample and average control EC50 at the given time point. Red shows a change towards resistance, while blue shows a change towards sensitivity. Samples are ranked along the x-axis from least-to-most sensitive. Sample labels on the x-axis are represented by darker colors the longer they have been evolved in the evolutionary experiment.

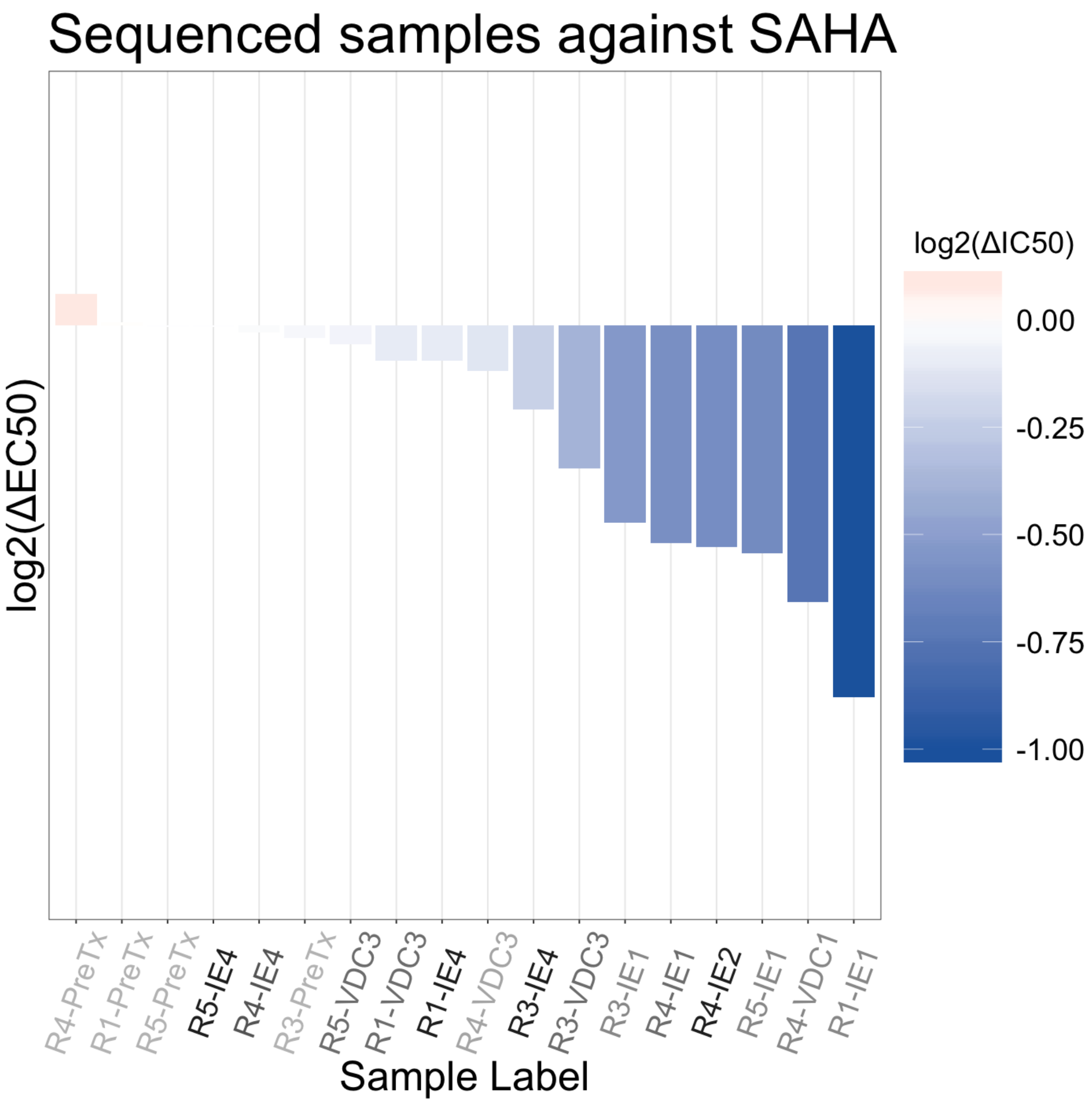


Supplementary Figure 12: **Waterfall of EC50 values for sequenced samples against vorinostat (SAHA), related to Figure 4.** Color represents $\log_2$ change in EC50 between the sample and average control EC50 at the given time point. Red shows a change towards resistance, while blue shows a change towards sensitivity. Samples are ranked along the x-axis from least-to-most sensitive. Sample labels on the x-axis are represented by darker colors the longer they have been evolved in the evolutionary experiment.

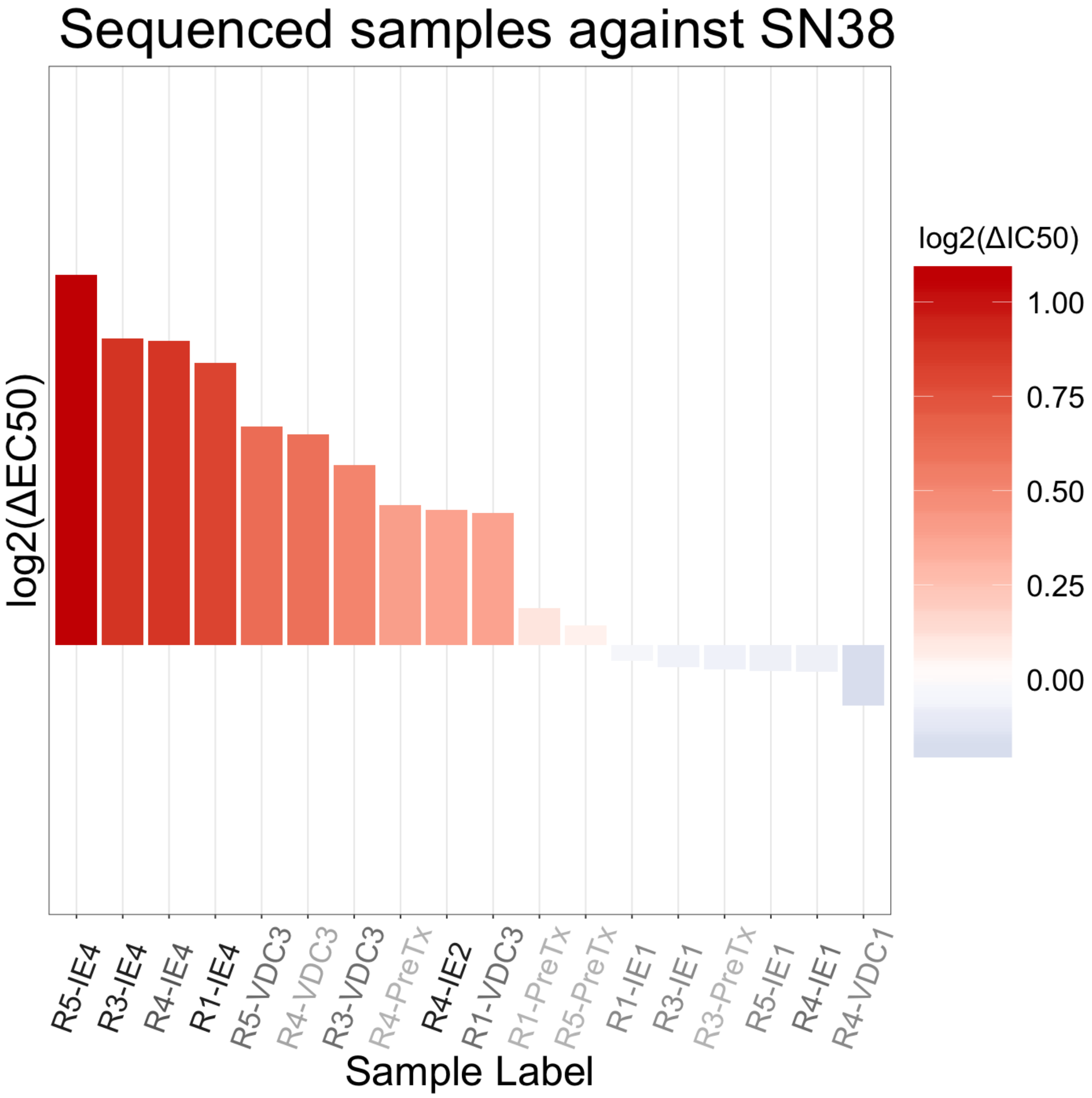


Supplementary Figure 13: **Waterfall of EC50 values for sequenced samples against irinotecan (active metabolite, SN38), related to Figure 4.** Color represents $\log_2$ change in EC50 between the sample and average control EC50 at the given time point. Red shows a change towards resistance, while blue shows a change towards sensitivity. Samples are ranked along the x-axis from least-to-most sensitive. Sample labels on the x-axis are represented by darker colors the longer they have been evolved in the evolutionary experiment.

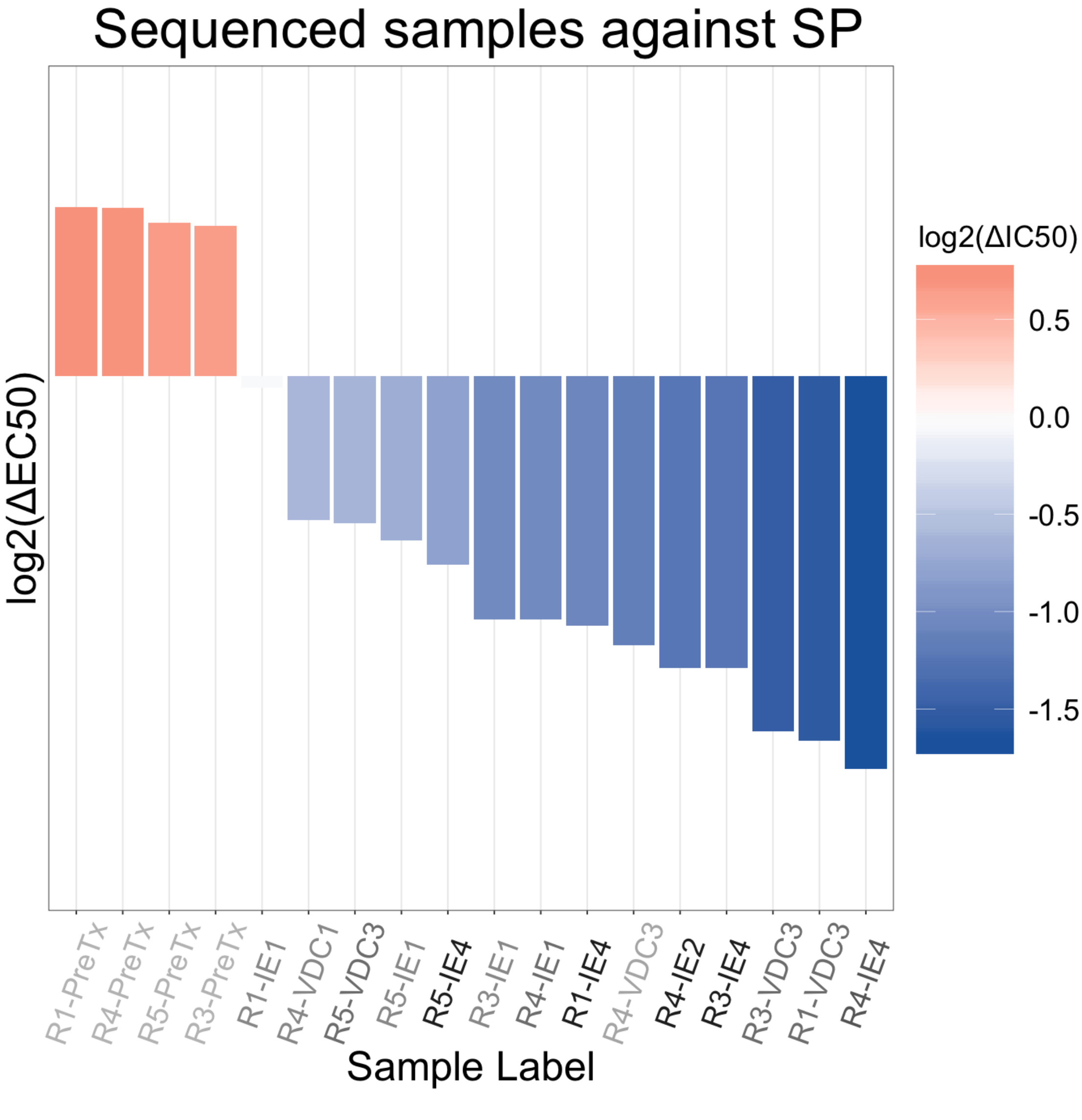


Supplementary Figure 14: **Waterfall of EC50 values for sequenced samples against SP-2509, related to Figure 4.** Color represents $\log_2$ change in EC50 between the sample and average control EC50 at the given time point. Red shows a change towards resistance, while blue shows a change towards sensitivity. Samples are ranked along the x-axis from least-to-most sensitive. Sample labels on the x-axis are represented by darker colors the longer they have been evolved in the evolutionary experiment.

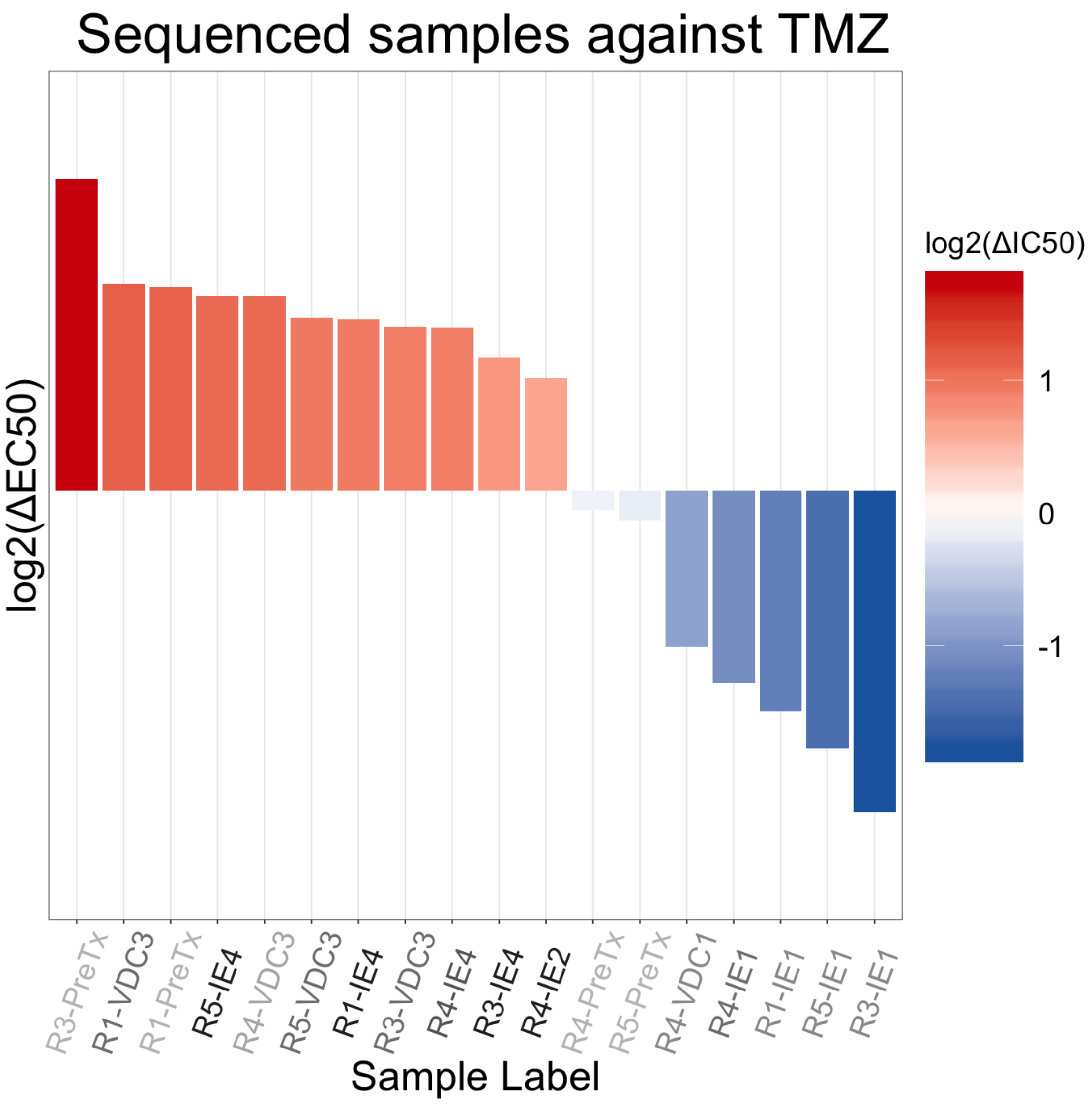


Supplementary Figure 15: **Waterfall of EC50 values for sequenced samples against temozolomide, related to Figure 4.** Color represents $\log_2$ change in EC50 between the sample and average control EC50 at the given time point. Red shows a change towards resistance, while blue shows a change towards sensitivity. Samples are ranked along the x-axis from least-to-most sensitive. Sample labels on the x-axis are represented by darker colors the longer they have been evolved in the evolutionary experiment.

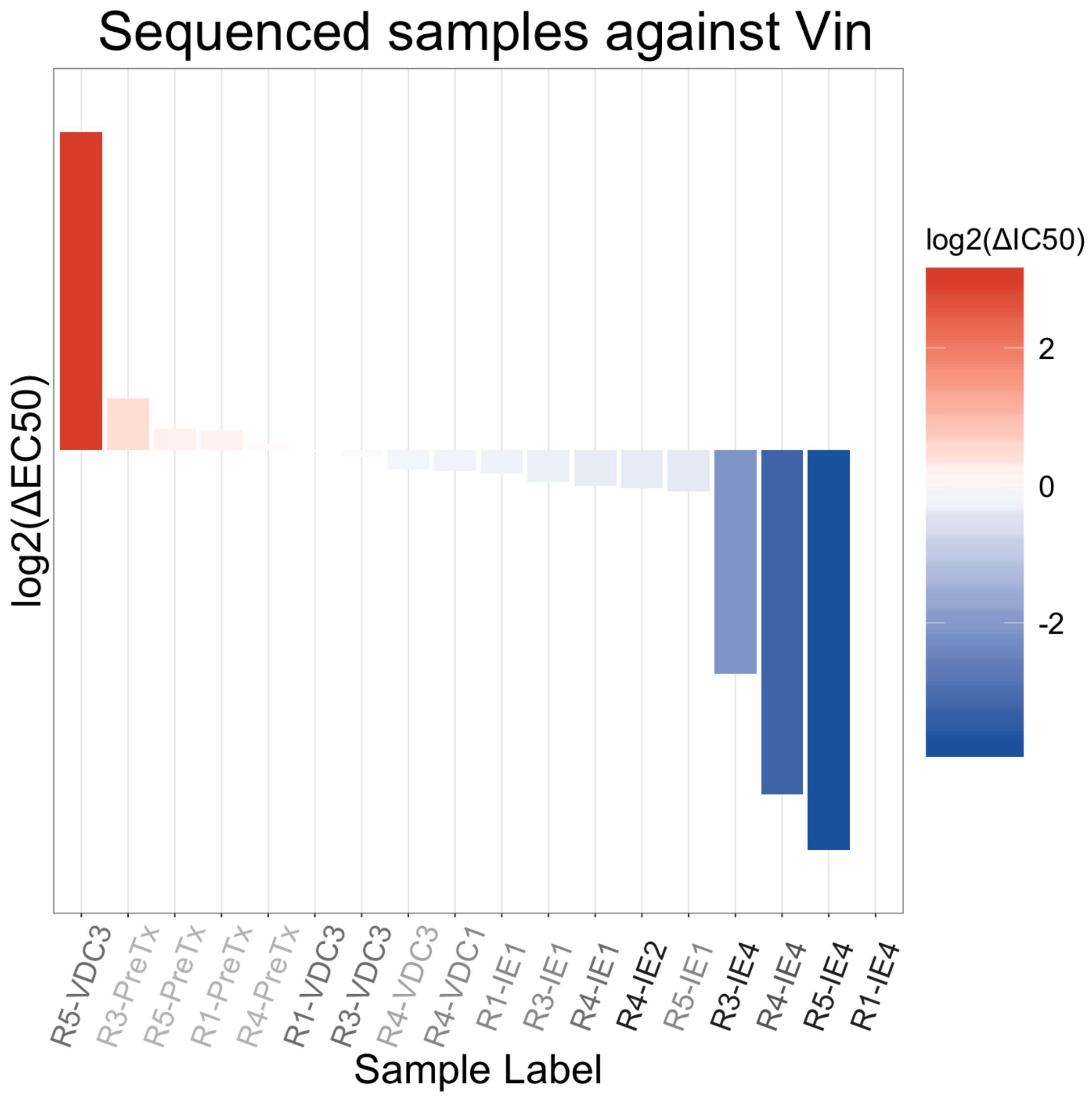


Supplementary Figure 16: **Waterfall of EC50 values for sequenced samples against vincristine, related to Figure 4.** Color represents $\log_2$ change in EC50 between the sample and average control EC50 at the given time point. Red shows a change towards resistance, while blue shows a change towards sensitivity. Samples are ranked along the x-axis from least-to-most sensitive. Sample labels on the x-axis are represented by darker colors the longer they have been evolved in the evolutionary experiment. The EC50 for the first replicate after the fourth exposure to the EC drug combination (R1-EC4) was indeterminant and removed from the waterfall plot.

### 2. Transparent Methods

*Materials*

EWS cells (A673 and TTC466 cells) were generous gifts from Dr. Stephen Lessnick at Nationwide Children's Hospital, Columbus, OH.

4-hydroperoxycyclophosphamide and sodium thiosulfate were purchased from Toronto Research Chemicals (North York, ON, Canada). Dactinomycin, SP-2509, doxorubicin, etoposide, temozolomide, pazopanib, olaparib, SAHA, and vincristine were products of Cayman Chemical (Ann Abor, MI). SN-38 was obtained from SelleckChem.com (Houston, TX). The classifications and abbreviations used for all these compounds are found in Table 1.

*Cell culture*

A673 cells were maintained in Dulbecco's Modified Eagle Medium (D-MEM) supplemented with 10% Fetal Bovine Serum (FBS) and penicillin and streptomycin at 37°C under humidified atmosphere containing 5% CO2. TTC466 cells were cultured in the same way except Roswell Park Memorial Institute (RPMI) medium was used instead of D-MEM.

*In vitro combination drug treatments to induce drug resistance*

Through drug toxicity assays, we determined EC50 concentrations of chemotherapeutics that are used as standard-of-care to treat EWS (Grier et al., 2003). This standard-of-care treatment for EWS consists of a vincristine-doxorubicin-cyclophosphamide (VDC) combination cycle followed by an etoposide-ifosfamide (EI) combination cycle. However, both cyclophosphamide and ifosfamide are prodrugs, requiring metabolic activation by an *in vivo* model. In the VDC drug combination, cyclophosphamide is replaced by 4-hydroxycyclophosphamide, an activated form of cyclophosphamide; however, there is no such commercially available option for ifosfamide. As ifosfamide is an analog of cyclophosphamide, it was also replaced by 4-hydroperoxycyclophosphamide, due to their similar chemical structures and mechanisms of action. Therefore, we recapitulate Ewing's sarcoma standard-of-care treatment regimen *in vitro* by cycling vincristine-doxorubicin-4-hydroxycyclophosphamide (VDC) and etoposide-4-hydroxycyclophosphamide (EC). The EC50 values were vincristine (0.8 and 0.9 nM), doxorubicin (0.015 and 0.023 nM), 4-hydroxycyclophosphamide (0.001 and 0.001 % by volume), and etoposide (0.7 and 0.37 $\mu$M) in the A673 and TTC466 cell lines, respectively.

In order to induce drug resistance in the A673 and TTC466 cell lines, each cell line was plated as 8 biological (evolutionary) replicates, where 5 experimental replicates were exposed to the drug combination cycles, described below, and 3 control replicates were maintained in dimethyl sulfoxide (DMSO). Due to contamination, one of the experimental replicates (Replicate 2) in the A673 cell line was discontinued. Experimental replicates were exposed to the standard-of-care drug cycles, as illustrated in Figure 1. Cells ($2 \times 10^6$ cells/10cm plate) were first exposed to a combination of vincristine, doxorubicin, and 4-hydroxycyclophosphamide at their EC50 concentrations. After 5 days of incubation, the medium was changed to maintenance medium without drugs. After they proliferate to sub-confluent density, a 10 cm plate was set for the next cycle with etoposide and 4-hydroxycyclophosphamide at their initial EC50 concentrations for 5 days, 96-well plates were set for drug sensitivity assay, and a fraction of cells were snap frozen for RNA extraction. Again, as the treated cultures grew to sub-confluence, this cycle was repeated with alternate exposure to the two drug combinations, along with drug sensitivity assays and sampling for RNA extraction between each drug cycle. On the fifth application of the VDC drug combination, the concentration of the drug combination was increased to 6 nM, 0.05 mM, and 0.006% by volume for vincristine, doxorubicin, and cyclophosphamide, respectively.

*Drug toxicity assay*

Cells were plated into 96-well plates at the density of 6,000 cells/90$\mu$l/well. The next day, 10 $\mu$l of medium containing various concentrations of drug of interest were added to each well. The final concentration of

DMSO used as solvent was kept constant (0.1% by volume for dactinomycin, SP2509, doxorubicin, etoposide, olaparib, SAHA, SN38, and vincristine; and 1% by volume for temozolomide and pazopanib).

4-hydroxycyclophosphamide was prepared freshly just prior to each assay by incubating 1 mg of 4-hydroperoxycyclophosphamide with 100 $\mu$L of water containing 1 mg sodium thiosulfate at room temperature for 30 sec, converting 4-hydroperoxycyclophosphamide to 4-hydroxycyclophosphamide. The resulting solution was used for toxicity assay starting with 0.2% (by volume) as the highest concentration. Matching dilution series of sodium thiosulfate solution was assessed as a control to assess 4-hydroxycyclophosphamide toxicity, again using 0.2% (by volume) as the highest concentration.

After five days of incubation, cell viability of each well was determined by measuring the enzymatic conversion of alamarBlue (Bio-Rad, Hercles, CA) (Hamid et al., 2004). After addition of alamarBlue solution (10 $\mu$l/well), the plate was incubated for two to four hours and the fluorescence intensity (excitation 560 nm / emission 590 nm) of each well was detected by Symphony H2(BioTek, Winooski, VT), a multi-well plate reader. Background fluorescence was determined by measuring the wells without cells incubated with alamarBlue.

*Drug response modeling and EC50 estimation*

Net alamarBlue conversion for each well was calculated by subtracting the average background fluorescence from each of the fluorescence values. A four-parameter log-logistic (LL.4) model (Hill function) was fit for each biological replicate, performed in triplicate, using the drm function from the drc package in R. This function models the survival measure $S(X)$ at a given dose $X$ as:

$$S(X) = b + \frac{a - b}{1 + \left(\frac{EC50}{X}\right)^H}$$

where $S(X)$ is the expected response at dose $X$, $a$ is the minimum response (when dose = 0), $b$ is the highest response (when dose = $\infty$), $EC50$ is the point of inflection (dose at which 50% of the response occurs), and $H$ (known as the Hill slope) is the steepest part of the curve (Gadagkar and Call, 2015). A negative value for H, as seen in these models, denotes a descending curve, while a positive $H$ represents an ascending curve. Estimated EC50 from these models was solved using the ED function from the drc package (version 3.0.1) in R.

Although there is no perfect measure of drug sensitivity and resistance, we chose EC50 as our measure of drug response, because of its compatibility with existing literature and because of the agreement of this model to experimental data. A caveat of this measure is the possibility for the maximum response, $b$ changes, while the point of inflection, $EC50$, for the curve does not (Jang et al., 2014). A plot of all dose-response triplicates with their estimated EC50 can be found in the linked GitHub repository.

*RNA extraction and sequencing*

Ribosomal-RNA depleted RNA was prepared from 18 samples of interest using RiboMinus Eukaryote Kit (ThermoFisher, Waltham, MA). RNA sequencing was performed at the Genomic Core, the Lerner Research Institute (Cleveland, OH) with HiSeq 2500 (Illumina, San Diego, CA). Quality control and read trimming was performed using fastp v0.20.0 (Chen et al., 2018). Read alignment was done using STAR v2.7.1 and alignment quantification was done using salmon v0.14.1 against gencode v31 transcript set with average 12 million reads per sample (Dobin et al., 2013; Patro et al., 2017; Harrow et al., 2012). Transcript level abundance estimates were then converted to gene level estimated counts using tximport R package (Soneson et al., 2015).

*Differential gene expression analysis*

Samples sent for sequencing were ranked based on their EC50 to each drug. For each drug analyzed, differential gene expression (DE) analysis compared samples in the top and bottom third of the ranked EC50 values. This DE analysis was performed using the EBSeq R package (version 1.24.0), with a false discovery threshold of 0.05 and the maxround parameter set to 15 (Leng and Kendziorski, 2019; Žiberna, 2018).

### References


Chen, S., Zhou, Y., Chen, Y., Gu, J., 2018. fastp: an ultra-fast all-in-one fastq preprocessor. Bioinformatics 34, i884–i890.

Dobin, A., Davis, C.A., Schlesinger, F., Drenkow, J., Zaleski, C., Jha, S., Batut, P., Chaisson, M., Gingeras, T.R., 2013. Star: ultrafast universal rna-seq aligner. Bioinformatics 29, 15–21.

Gadagkar, S.R., Call, G.B., 2015. Computational tools for fitting the hill equation to dose–response curves. Journal of Pharmacological and Toxicological methods 71, 68–76.

Grier, H.E., Krailo, M.D., Tarbell, N.J., Link, M.P., Fryer, C.J., Pritchard, D.J., Gebhardt, M.C., Dickman, P.S., Perlman, E.J., Meyers, P.A., et al., 2003. Addition of ifosfamide and etoposide to standard chemotherapy for ewing's sarcoma and primitive neuroectodermal tumor of bone. New England Journal of Medicine 348, 694–701.

Hamid, R., Rotshteyn, Y., Rabadi, L., Parikh, R., Bullock, P., 2004. Comparison of alamar blue and mtt assays for high through-put screening. Toxicology in vitro 18, 703–710.

Harrow, J., Frankish, A., Gonzalez, J.M., Tapanari, E., Diekhans, M., Kokocinski, F., Aken, B.L., Barrell, D., Zadissa, A., Searle, S., et al., 2012. Gencode: the reference human genome annotation for the encode project. Genome research 22, 1760–1774.

Jang, I.S., Neto, E.C., Guinney, J., Friend, S.H., Margolin, A.A., 2014. Systematic assessment of analytical methods for drug sensitivity prediction from cancer cell line data, in: Biocomputing 2014. World Scientific, pp. 63–74.

Leng, N., Kendziorski, C., 2019. EBSeq: An R package for gene and isoform differential expression analysis of RNA-seq data. R package version 1.24.0.

Patro, R., Duggal, G., Love, M.I., Irizarry, R.A., Kingsford, C., 2017. Salmon provides fast and bias-aware quantification of transcript expression. Nature methods 14, 417.

Soneson, C., Love, M.I., Robinson, M.D., 2015. Differential analyses for rna-seq: transcript-level estimates improve gene-level inferences. F1000Research 4.

Žiberna, A., 2018. Generalized and Classical Blockmodeling of Valued Networks. R package version 0.3.4.
